# The *Aedes aegypti* bacterial microbiota is robust to infection with the obligate microsporidian parasite *Edhazardia aedis*

**DOI:** 10.64898/2026.01.28.702404

**Authors:** Dom Magistrado, Sarah M. Short

## Abstract

*Edhazardia aedis* is an obligate microsporidian parasite of the arthropod vector *Aedes aegypti*, which is responsible for the spread of several vertebrate pathogens of global health importance. *E. aedis* can be highly virulent to *Ae. aegypti* and infection has severely detrimental effects on multiple life history traits that are relevant to the vectoral capacity of *Ae. aegypti*, including longevity, body size, propensity to host-seek and blood-feed, and reproductive capacity. Because *E. aedis* is also highly specific to *Ae. aegypti* and is incapable of completing its full life cycle in any other mosquito species, *E. aedis* merits investigation as a novel tool for biological vector control. In the present study, we queried the effect of *E. aedis* infection on the bacterial microbiota of adult female *Ae. aegypti* using high-throughput amplicon sequencing of the 16S rRNA gene. Analysis of sequencing data revealed that the bacterial microbiota community is strikingly robust to *E. aedis* infection, as we observed no significant effect on alpha or beta diversity, differential abundance of any taxa, predicted metabolic function profile, or overall bacterial load. The data show that *E. aedis*, despite dramatically impacting the health and fitness of the adult female mosquito, does not affect the microbiota. These results provide unique insight into tripartite relationships (or lack thereof) between hosts, pathogens, and the microbiota.

## Introduction

Microbes that live in association with host organisms, i.e. the microbiota, are increasingly implicated in a holistic understanding of the biology of the host. The holobiont concept is the idea that host organisms are best understood not in isolation, but rather, as a complex or “meta-organism” comprised of both the host and the microbes associated with that host (1,2). The microbiota affect a suite of functions in hosts, playing key roles in immunity and metabolism (3–5), and perturbation of the microbiota (dysbiosis) typically causes disease in hosts (6–8). Interest in understanding the mosquito microbiota has emerged in recent years (9–13) because microbes can be exploited to suppress the spread of mosquito-borne diseases such as dengue, Zika, chikungunya, and yellow fever. Indeed, microbes such as the endosymbiotic bacteria *Wolbachia* and the entomopathogen *Bacillus thuringiensis israelensis* have been utilized to suppress vector populations with some success (14,15). Despite these successes, mosquito-borne disease remains a growing global public health problem, as 2024 was the worst year for dengue cases recorded in history (16). Clearly, the demand for new solutions remains high. Studying mosquito-microbiota interactions provides not only potential for uncovering novel microbial agents with direct potential for biocontrol, but also contributes to an improved, holistic framework of understanding mosquito vector physiology that accounts for the increasingly realized fundamental role of microbes.

Mosquitoes associate with microbes at every life stage (17) and the microbiota play important roles in many physiological processes. For example, the microbiota are indispensable for normal larval development (18), may mediate insecticide resistance (19– 23), play a role in the formation of the adult peritrophic matrix (24), and contribute to the digestion of both blood (17,24,25) and sugar (26–29). Female mosquitoes may even beneficially engineer the microbiota of their progeny by depositing microbes into the larval habitat during oviposition (30). While the mosquito microbiota is taxonomically diverse and includes bacteria, viruses, fungi, and protists, bacteria are by far the best studied and have disproportionately informed the current paradigm of how resident microbiota interact with hosts (10). Importantly, all microbes within this diverse suite have the potential to interact not only with the mosquito directly, but with other members of the holobiont as well, and this concept underpins the field of microbial ecology. Indeed, exploration of tripartite interactions between mosquitoes, resident microbiota, and microbial pathogens of human importance have revealed that the mosquito’s resident bacterial, viral, fungal, and microsporidian microbiota can interfere with the success of *Plasmodium* parasites and multiple arboviruses including dengue and West Nile in establishing infection in mosquitoes (31–42).

In the present study, we investigate the impact of the microsporidian parasite *Edhazardia aedis* on the bacterial microbiota of the yellow fever mosquito *Aedes aegypti*. Microsporidia are fungus-like obligate intracellular parasites that are strikingly ubiquitous and diverse, infecting myriad hosts ranging from protists to mammals (43), including hosts from all major insect orders (44). *E. aedis* was first isolated in Puerto Rico in 1930, then later re-discovered in Thailand in 1979 (45–47). The current range of *E. aedis* has not been surveyed and thus remains unknown. Naïve *Ae. aegypti* larvae can encounter *E. aedis* in the aquatic larval habitat through ingestion of spores released from infectious larvae (i.e., horizontal infection). After spores enter the larval digestive tract, they invade cells and aggregate in the gastric caecae (48). *E. aedis* is virulent to *Ae. aegypti*, and survival outcome is affected by larval instar development stage, food availability, and the quantity of spores ingested (49,50). In females that survive to adulthood, *E. aedis* is limited primarily to the oenocytes of the fat body until bloodfeeding, an event that initiates infection of the ovaries and infection of progeny, thereby potentiating transovarial vertical transmission of the parasite (48). *E. aedis* undergoes development and replication in the vertically infected larvae, leading to the formation of distinctive spore cysts that, upon larval death, are released into the larval habitat to permit horizontal infection of naïve larvae, thus completing the life cycle (48). Infection with *E. aedis* alters the whole-body transcript abundance of genes related to immunity and metabolism at all life stages in horizontally infected female mosquitoes (51). It also reduces defense against opportunistic bacterial infection (52) and begets high fitness costs to the mosquito host via reductions in longevity, body size, propensity to host-seek, propensity to blood feed, and reproductive capacity (47,53–55). Importantly, *E. aedis* exhibits a high degree of host specificity to *Ae. aegypti*; while some other mosquito species can become infected with *E. aedis* via horizontal transmission, vertical transmission has not been recorded in any species other than *Ae. aegypti* (56,57). Due to *E. aedis’s* high vertical transmission fidelity to *Ae. aegypti*, its mosquitocidal properties, and its effects on life history traits that are relevant to vectorial capacity, *E. aedis* warrants exploration as a strategy for controlling *Ae. aegypti* populations. Because understanding the effects of biological control agents on non-target microbial communities is important (58–60), and because infection with diverse pathogens including microsporidians has been shown to affect the arthropod microbiota in numerous instances (61–72), we sought to assess the effects of *E. aedis* infection on the microbiota of *Ae. aegypti*.

In the present cross-sectional study, we query the effects of horizontal infection with *E. aedis* on the abdominal bacterial microbiota of individual adult female *Ae. aegypti* (non-blood-fed). We use high throughput amplicon sequencing targeting the 16S rRNA gene followed by ecological analyses suitable for the compositional nature of sequencing data (73) to capture infection-induced perturbations in bacterial diversity. Because sequencing counts do not reflect the absolute abundance of starting material, we also use quantitative PCR targeting the 16S rRNA gene to approximate, and query an effect of *E. aedis* infection on, bacterial load. Overall, the data indicate that the bacterial microbiota is strikingly robust to *E. aedis* infection, as no analyses revealed any effect of infection on abundance, taxonomic diversity, or predicted metabolic functional profiles of the microbial communities. These findings contribute novel information about the nature and specificity of *Ae. aegypti’s* physiological response to *E. aedis* infection and provide an improved understanding of the mosquito holobiont.

## Results

To query how horizontal infection with *Edhazardia aedis* impacts the bacterial microbiota of *Aedes aegypti*, we reared three paired sets of *E. aedis*(+) or mock-infected larvae in parallel from egg to adulthood and extracted DNA from n=7-8 individual female mosquitoes (non-blood-fed) per cage shortly after eclosion (Fig. 1). To characterize both the abundance and the composition of the bacterial microbiota, we quantified and sequenced a segment of the V3-V4 hypervariable region of the bacterial 16S rRNA gene using quantitative PCR and high-throughput sequencing.

**Figure 1.**
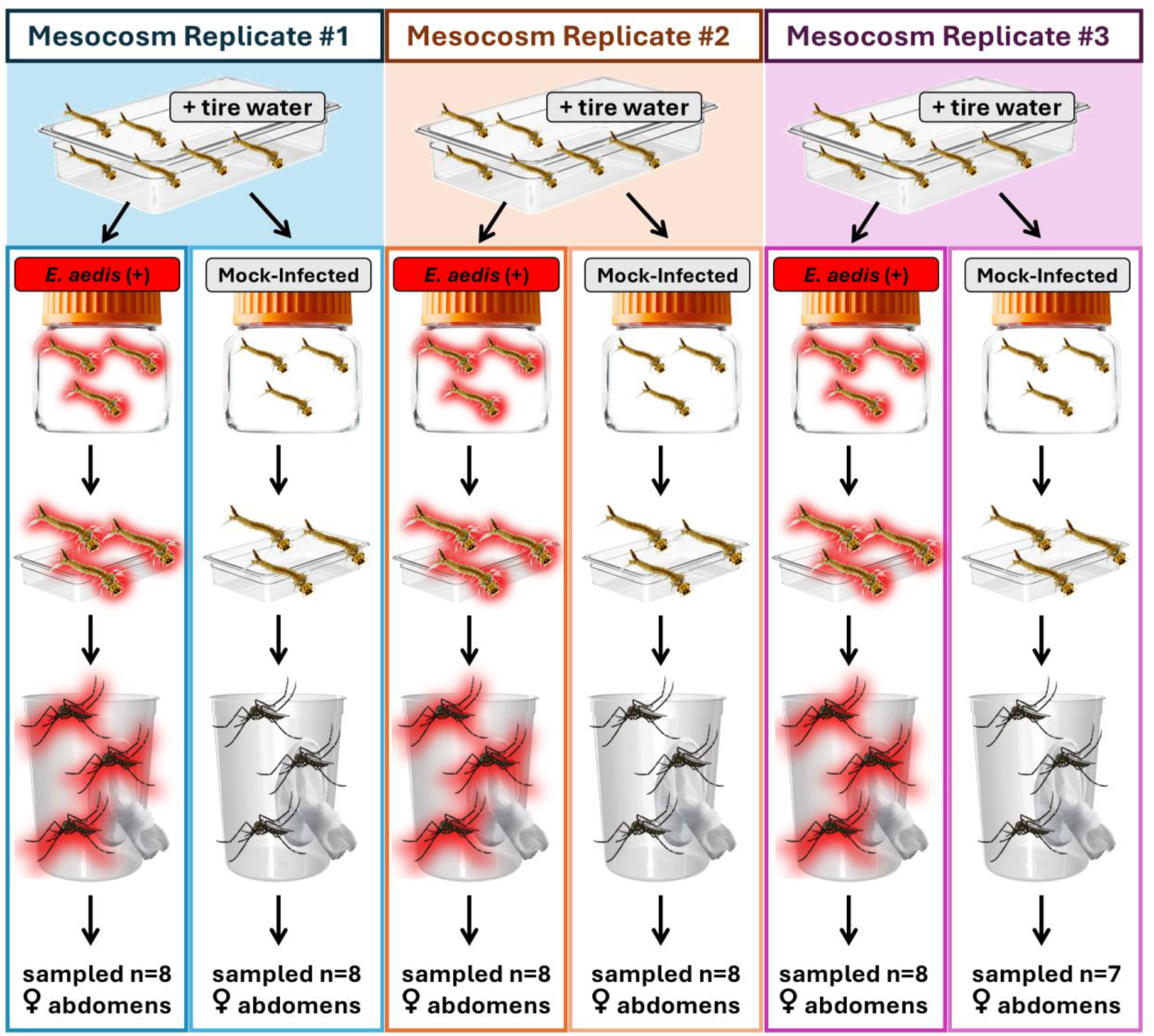
Experimental Design Schematic. One egg paper was used to hatch all *Ae. aegypti* (LVP strain) mosquitoes used in this experiment. Immediately after hatching, larvae were divided into three trays. In an effort to more closely mimic natural larval environments as opposed to standard laboratory rearing, each tray was inoculated with water collected from *Aedes* larval development sites. Larvae were reared in trays until they reached instar 2-3, then were designated to a control (mock-infected) or *E. aedis*(+) group. Groups destined for *E. aedis* infection were horizontally infected via exposure to purified *E. aedis* spores harvested from vertically infected larvae (not shown). Following horizontal infection, larvae were transferred to standard rearing trays and monitored until pupation. Pupae were transferred to cages and allowed to eclose. At 3-5 days post-eclosion, n=7-8 abdomens from individual females were sampled from each cage and subjected to DNA extraction before sequencing.

### Characterization of sequencing dataset

The final sequencing dataset used for analysis contained 5,757,955 total reads with an average of 122,510 reads per sample, and 3,667 total ASVs with an average of 221 ASVs per sample. Rarefaction curves generated from the final phyloseq object suggest that the samples were sequenced to a sufficient depth (Fig. S1). Twenty “Core ASVs” (0.01% of all ASVs) were present in all 47 samples (Table S1).

### Taxonomic Composition

The bacterial microbiota was dominated primarily by bacteria from the phyla *Pseudomonadata* and *Bacillota*, with a substantial proportion of reads also coming from the phylum *Actinomycetota* (Fig. 2). Within *Pseudomonadata, Gammaproteobacteria* were more abundant than *Alphaproteobacteria. Bacilli* was the dominant class within *Bacillota* and *Actinobacteria* was the dominant class within *Actinomycetota*.

**Figure 2.**
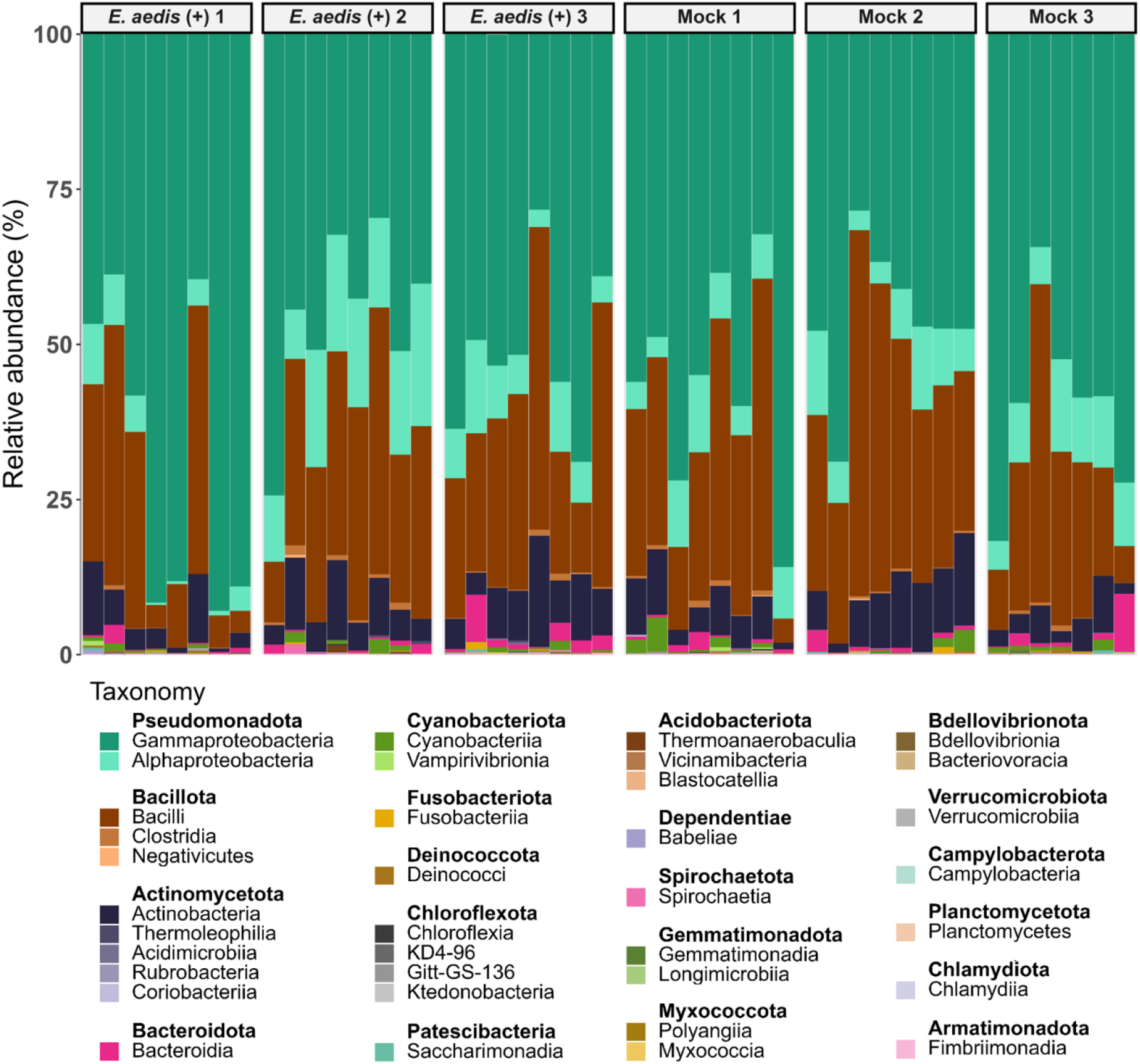
The bacterial microbiota is dominated by *Pseudomonadata, Bacillota*, and *Actinomycetota*. Individual bars represent individual samples of n=1 female *Ae. aegypti* abdomen.

Sequencing of the tire water that was used to inoculate all larval trays revealed that out of the 286 ASVs present in the tire water, 132 ASVs were also detected in mosquito samples (“Shared” ASVs), while 163 ASVs were unique to the tire water inoculum (Fig. 3). Shared ASVs comprised approximately 50.0% of the tire water reads (90,950 out of 182,007 reads) and 57.1% (3,288,839 out of 5,757,955 reads) of the mosquito sample reads. Shared ASVs constitute a majority (16 out of 20) of the Core ASVs. At the phylum level, the composition of the Shared ASVs is similar to that of the ASVs in mosquito samples only, comprising mostly *Pseudomonadata* and *Bacillota*, with a smaller proportion of *Actinomycetota*.

**Figure 3.**
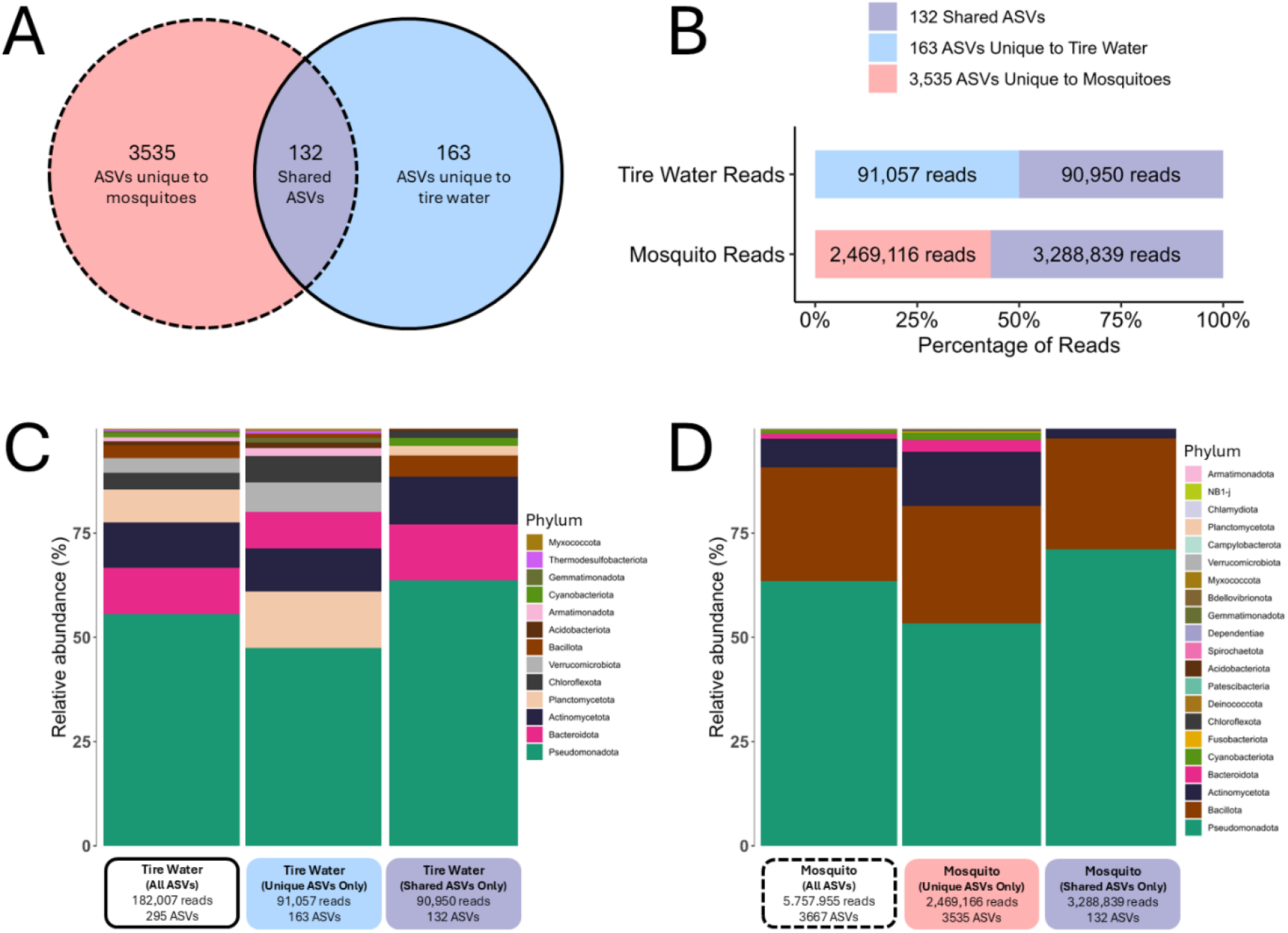
Analysis of ASVs in tire water. ASVs were A) categorized as unique to mosquito samples, unique to tire water inoculum, or shared between the two sample types, and B) the weights of each categorization were characterized using the number or proportion of reads assigned to each categorization within tire water or mosquito reads. Phylum-level taxonomy is displayed for tire water (C) or mosquito samples (D) as uncategorized (left bar plots) or categorized into unique or shared ASVs (middle and right bar plots, respectively).

### Infection with Edhazardia aedis does not alter the abundance, taxonomic diversity, or predicted metabolic functional profiles of the bacterial microbiota in female non-blood-fed Aedes aegypti

We characterized the alpha diversity of the bacterial microbiota in each sample using Chao1, Shannon, and Simpson indices and queried the effects of *E. aedis* infection and larval mesocosm replicate on each alpha diversity index using the Wilcoxon Rank-Sum test (for *E. aedis* infection status) or the Kruskal-Wallis test (for larval mesocosm replicate). Neither *E. aedis* infection status nor mesocosm had a significant effect on any measure of alpha diversity of the adult bacterial microbiota (Fig. 4; File S1: Wilcoxon rank-sum test: p_Chao1, *E. aedis*_=0.246, p_Shannon, *E. aedis*_=0.907, p_Simpson, *E. aedis*_=0.742; File S1: Kruskal-Wallis test: p_Chao1, Mesocosm_=0.634, p_Shannon, Mesocosm_=0.161, p_Simpson, Mesocosm_=0.126)).

**Figure 4.**
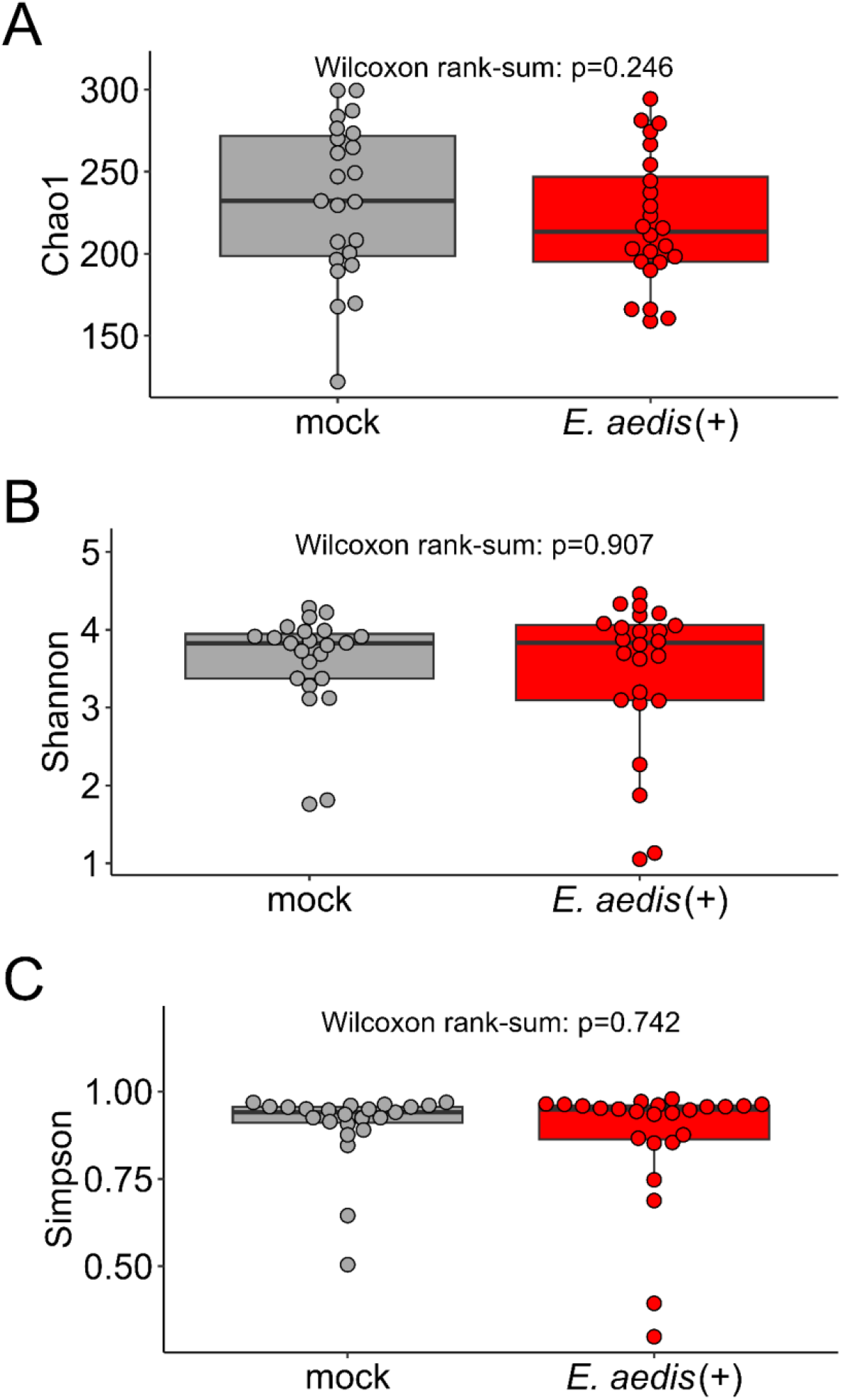
Infection with *E. aedis* does not alter the alpha diversity of the bacterial microbiota in the abdomens of *Ae. aegypti*. P-values indicate the results of Wilcoxon rank-sum statistical tests for significant differences in A) Chao1 Index, B) Shannon Index, or C) Simpson Index between mock-infected and *E. aedis*(+) samples. No differences were found at any taxonomic level; ASV level is shown in the figure.

We constructed beta diversity ordinations using a suite of distances or transformed abundance matrices: Bray Curtis distances (phylogeny-unaware), robust CLR transformed abundance matrix (phylogeny-unaware, appropriate for compositional data), CLR transformed abundance matrix with imputed zeroes (phylogeny-unaware, appropriate for compositional data), and balances derived from a phylogenetic isometric log-ratio transformation (phylogeny-aware, appropriate for compositional data) (Fig. 5). To query effects of *E. aedis* infection on beta diversity of the bacterial microbiota, we used PERMANOVAs to test for significant differences between *E. aedis*(+) and mock-infected groups, with restricted permutations to account for any effect explained by larval mesocosm replicate. PERMANOVAs revealed no significant effect of infection on beta diversity of the bacterial microbiota for any distance metric (Fig. 5; File S1: PERMANOVAs: p_Bray-Curtis, *E. aedis*_=0.368, p_RCLR, *E. aedis*_=0.698, p_Zero-imputed CLR, *E. aedis*_=0.245, p_PHILR, *E. aedis*_=0.428).

**Figure 5.**
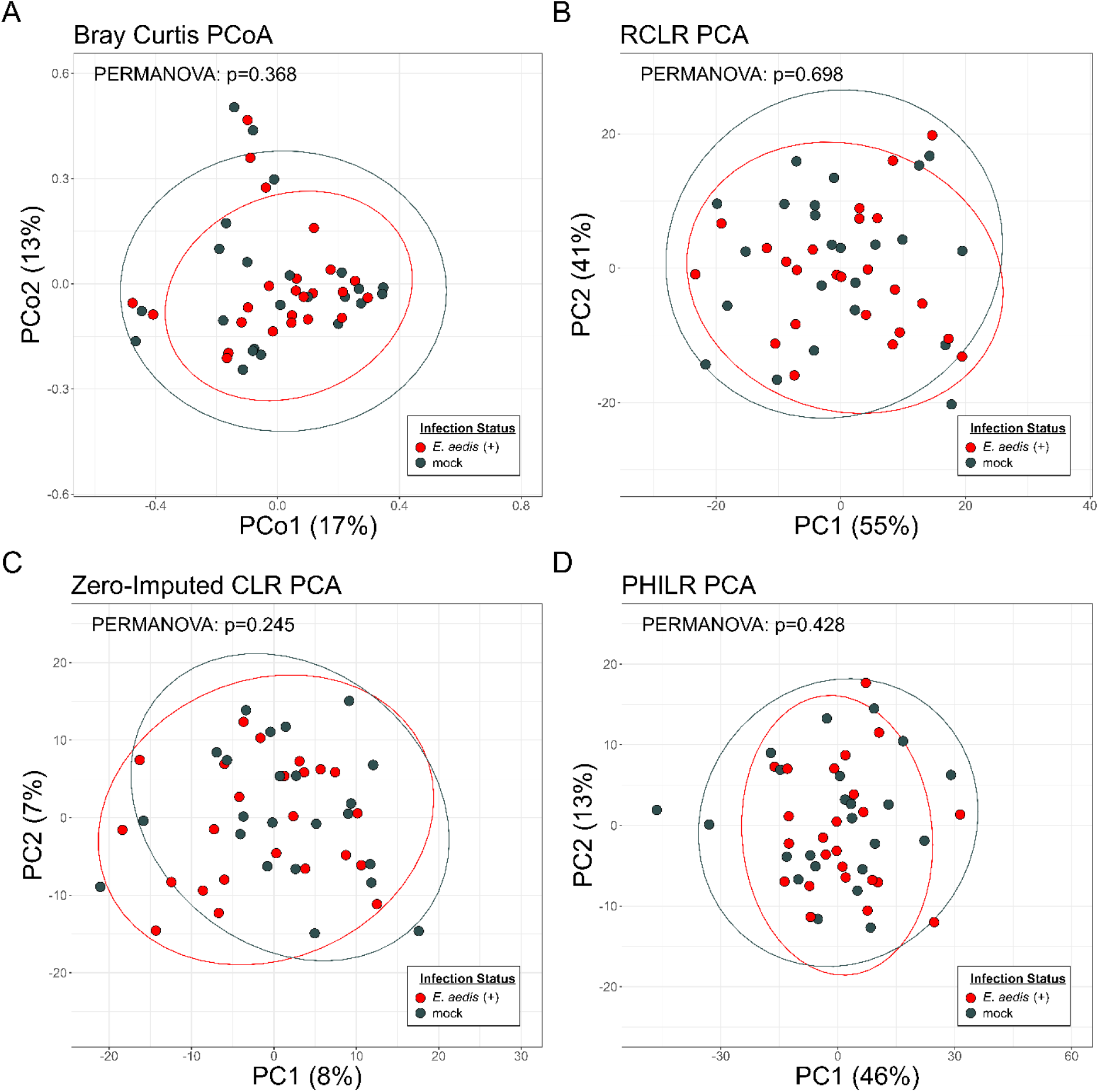
Infection with *E. aedis* does not alter the beta diversity of the bacterial microbiota in the abdomens of *Ae. aegypti*. Ordination analyses use A) Bray-Curtis distances, B) Robust CLR distances, C) zero-imputed CLR distances, or D) PHILR distances (phylogeny-aware). P-values indicate the results of PERMANOVA tests for significant differences between mock-infected and *E. aedis*(+) samples. No differences were found at any taxonomic level; ASV level is shown in the figure.

We also sought to understand the effect of *E. aedis* infection on bacterial load. Because sequencing data cannot reflect the absolute abundance of starting material, we performed quantitative PCR targeting the 16S rRNA gene to estimate the bacterial load associated with each sample. Mixed effects linear regression analysis of qPCR data revealed no effect of *E. aedis* infection on bacterial load (Fig. 6; File S1: p_*E. aedis*_=0.819).

**Figure 6.**
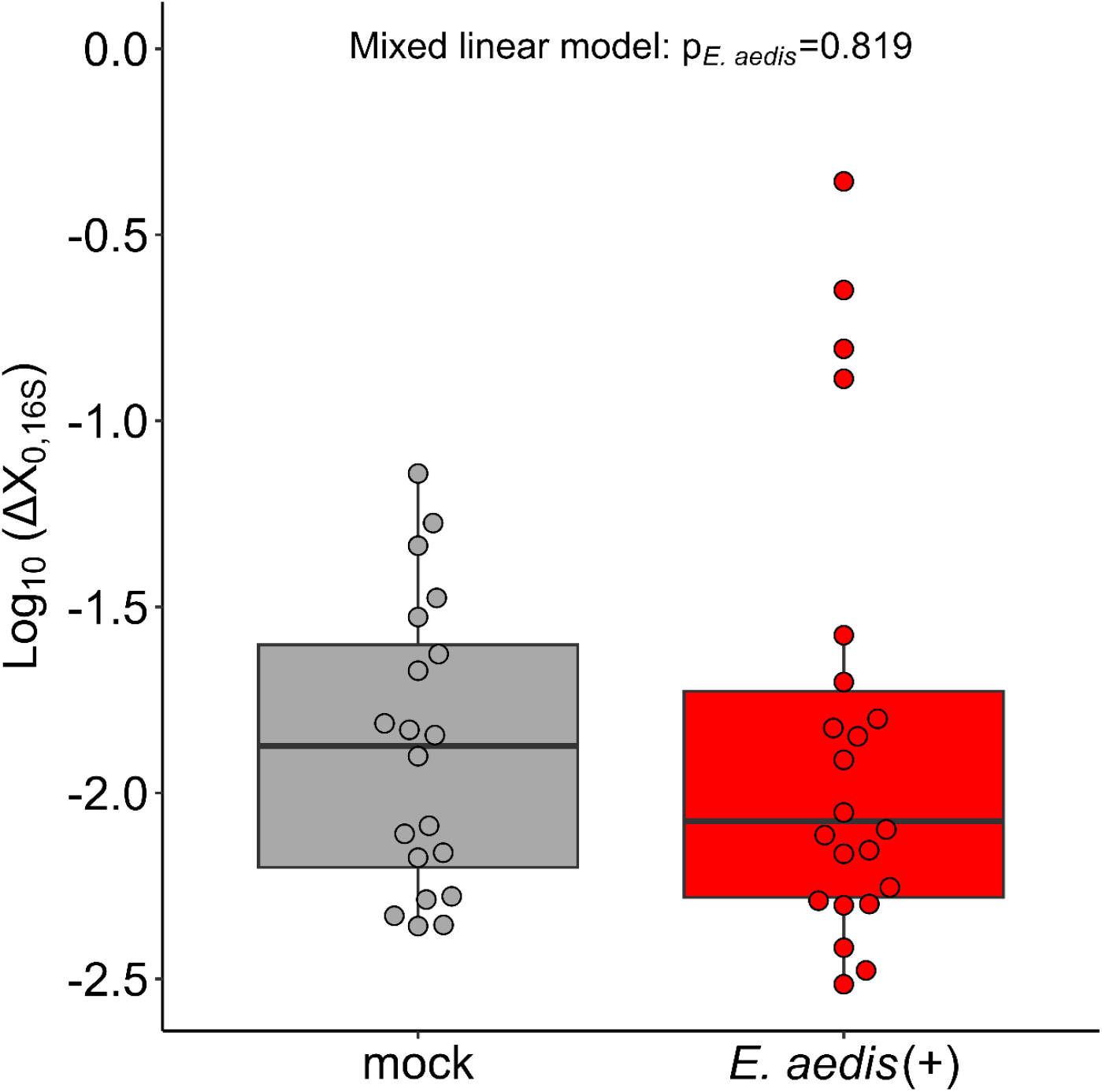
Infection with *E. aedis* does not alter bacterial load in the abdomens of *Ae. aegypti*. The y-axis displays log10-transformed ΔX_0,16S_ values, which were obtained from analysis of quantitative PCR data, and represent the starting amount of 16S bacterial DNA in each sample normalized to the amount of *RpS7* reference mosquito DNA. ΔX_0,16S_ values serve as a proxy for measurement of bacterial load. The p-value indicates the result of a mixed linear model testing the effect of *E. aedis* infection on log10-transformed ΔX_0,16S_ values.

We used three differential abundance analyses to determine differential abundance of taxa between Infection Status groups: ANCOM-BC2, ALDeX2, and LinDA. Overall, differential abundance analysis did not reliably identify any taxa that were consistently differentially abundant in infected mosquitoes compared to mock-infected. Finally, functional prediction analyses FAPROTAX and Tax4Fun2 revealed no effect of infection on the metabolic functional profiles of the bacterial microbiota.

## Discussion

Our data show that the non-blood-fed female *Ae. aegypti* abdominal bacterial microbiota is robust to horizontal infection with the microsporidian parasite *E. aedis*. In contrast, a diverse set of studies conducted in *Aedes* and *Anopheles* mosquitoes show that infection with arboviruses, malaria parasites, or entomopathogenic fungi, perturb the mosquito’s resident microbial community structure in various ways (69,72,74–76). Similar effects have also been observed in a diversity of microsporidian-arthropod associations, including honey bees, silkworms, and Pacific white shrimp (61,63,65–68,71). A long-standing topic of interest within this context is the mechanism by which such perturbations occur (72). Specifically, perturbation of the microbiota may result from direct microbial interactions between specific taxa or occur secondarily to an immune response mounted against the invading microbe (77). Our study adds novel context to this topic: unlike most of the pathogens referenced in studies above, *E. aedis* never encounters the gut lumen or invades gut cells in adult mosquitoes and, as a result, cannot directly interface with the gut microbiota. Interestingly, the adult mosquito does mount a marked transcriptional response to *E. aedis* infection that includes several immune signaling genes, although tissue specificity of this response has not yet been studied (51). *E. aedis* infection therefore provides a unique natural infection system for uncoupling the potential for direct microbe-microbe interactions within the mosquito gut from indirect interactions occurring only as a result of infection on the host.

Interestingly, we previously showed that female, non-blood-fed *E. aedis*-positive *Ae. aegypti* orally exposed to *Serratia marcescens* displayed higher *S. marcescens* loads compared to *E. aedis*-negative mosquitoes (52). Notably, *S. marcescens*, which is a common microbiota member and opportunistic pathogen, was not detected in the present study. Taken together with the findings in the present study, the data indicate that *E. aedis* can modulate the relationship between the host and gut microbes, but the effect is context-dependent. Two hypotheses emerge: 1) It is possible that *S. marcescens* is, unlike other microbiota members, uniquely capable of exploiting *E. aedis* infection-induced disease in the mosquito to establish and proliferate. Under such a scenario, the results of the present study may have differed had *S. marcescens* been present in the microbiota. 2) Another explanation is that *E. aedis* alters the mosquito’s induced physiological response resulting from acute bacterial exposure (e.g., oral delivery of *S. marcescens*), but not the host mechanisms underlying the maintenance of a diverse resident microbiota community.

Importantly, more work distinguishing the mechanisms by which the mosquito host responds to invading microbes versus those by which the mosquito host shapes the resident gut microbiota is needed to integrate these findings. It is well-established that the diversity of the microbiota is strongly influenced by both exogenous conditions (e.g., nutrition, habitat) and endogenous host factors (e.g., taxonomy, sex, development stage), but only a limited number of studies have explored the mechanistic underpinnings of this governance (78). More broadly, an emerging question in biology is how and why hosts may (or may not) shape the microbiota in response to stimuli such as infection, immune challenge, or disease (79–83). Some evidence in the literature indicates that processes outside of the gut can influence the composition of the gut microbiota. For example, wing necrosis in *Drosophila melanogaster* leads to the overgrowth of a gut commensal, although the signaling mechanism remains unclear (84). In the social ant species *Temnothorax rugatulus*, presenting a cuticular immune challenge (lipopolysaccharide and peptidoglycan) leads to immune system signaling-mediated decreases in gut microbiota diversity and overgrowth of bacteria from the genera *Entomoplasma* and *Pseudomonas* (85). These studies indicate that processes shaping the microbiota are not limited to the tissues colonized by the microbiota in some arthropods. In more complex and well-studied taxa, the opportunistically pathogenic nature of the microbiota is increasingly realized, and there are many instances of both infectious and non-infectious disease resulting in dysbiosis (86,87). Opportunistically pathogenic microbiota members can exploit pre-existing host disease to establish opportunistic infections (88). *E. aedis* is highly virulent to *Ae. aegypti*, and substantial work shows that infection is associated with disease (47,53– 55). However, our data show that the microbiota do not exploit the mosquito’s disease state and rather, remain unchanged. Possibly, *E. aedis* engineers host signaling to prevent the microbiota from detecting the host’s disease. Such a strategy would benefit *E. aedis* by preventing other microbes from, for example, killing the host before *E. aedis* can complete its life cycle, further activating the immune response in a manner that is deleterious to *E. aedis*, or competing with *E. aedis* for host-provisioned resources.

In a recent study, the presence of microsporidia in field-caught mosquitoes in Poland was positively correlated with the presence of two bacteria belonging to *Spiroplasmataceae* and *Leuconostocaceae* in some, but not all mosquito species represented in the sample (89). Furthermore, microsporidia infection was associated with a significantly greater proportion of predicted metabolic function dedicated to biosynthesis of ansamycins, the biosynthesis of vancomycin group antibiotics, and the pentose phosphate pathway (89). Neither *Ae. aegypti* nor *E. aedis* were represented in the study. Consistent with our study however, Trzebny et al. did not identify any effects of microsporidia infection on alpha or beta diversity of the bacterial microbiota, suggesting that most mosquito-associated microsporidia do not cause overall microbial community shifts in hosts. It is possible that microsporidians only interact with specific microbial taxa (e.g., *Spiroplasmataceae* and *Leuconostocaceae*) that were not present in the microbiota associated with our mosquitoes. Alternatively, aspects of microsporidia biology or pathogenesis that are unique to certain taxa may underlie differing effects of infection on the host microbiota (e.g. idiosyncratic tissue specificity or mode of infection). Furthermore, while the mosquitoes used in our study were non-blood-fed, field-collected mosquitoes may have blood-fed one or multiple times. Multiple mosquito-associated microsporidian species undergo important life cycle progression in response to blood feeding, including *E. aedis, Amblyospora connecticus*, and *Microsporidia MB* (44,90). Because the blood meal is inextricably connected to *Ae. aegypti* reproduction and immunity (91–96) and transiently but dramatically reshapes the diversity of the gut microbiota (97–99), investigating an effect of *E. aedis* infection in blood-fed mosquitoes (in which *E. aedis* has colonized the ovaries) would be of interest.

## Conclusion

The data indicate that the *Ae. aegypti* bacterial microbiota is unaltered in response to infection with *Edhazardia aedis*, a microsporidian pathogen with potential for biological control of *Aedes aegypti* mosquitoes. Because effective biocontrol agents should minimize effects on non-target microbial communities (58–60), the results of this study support the continued investigation of *E. aedis* as a strategy for controlling *Ae. aegypti* populations, although further research is needed (e.g. repeating the study at different stages of *E. aedis* infection). Interestingly, the absence of an effect of infection on the indigenous microbiota in this system contrasts with a multitude of diverse arthropod-pathogen systems in which the microbiota is altered by pathogenic infection, including microsporidian pathogens (61– 72). However, our findings are consistent with a prior study in mosquitoes that also revealed no impact of microsporidia infection on mosquito bacterial community structure (89). Additional research is needed to uncover the mechanisms underlying host regulation of the microbiota and how such regulation may or may not intersect with infection-induced changes in host physiology (e.g. immunity, metabolism). Overall, the present study motivates continued investigation of *E. aedis* as a biocontrol agent and provides insight into tripartite interactions (or a lack thereof) between vector mosquitoes, pathogens, and the microbiota.

## Materials and Methods

### Mosquito Maintenance

Throughout the experiment, all mosquitoes described in current and subsequent sections were maintained in a 27 °C chamber with 80% relative humidity under a 14 h:10 h light:dark cycle.

### Edhazardia aedis spore isolation and purification

*Edhazardia aedis* spores were harvested from vertically infected *Aedes aegypti* (LVP strain) as described in Grigsby et al., 2020 (100). To obtain spores for horizontal infection, eggs laid by *E. aedis*-infected females were hatched in RO water in a vacuum chamber. Upon hatching, larvae were thinned into 12.75” x 10.44” x 2.5” polycarbonate trays (Cambro, Huntington Beach, CA) containing RO water at a density of 100 individuals per tray. At this time, each tray was given a pinch of fish food (Tetramin) and half of a green cat pellet (9 Lives). Larvae were food restricted, but not starved, only for the purpose of extending time to pupation and permitting enough time for the development of binucleate *E. aedis* spores (100).

At day 10 post-hatch, 10 larvae were collected, washed with DI (deionized) water, and homogenized. Each individual homogenate was visually examined under a microscope and evaluated for the presence of binucleate spores. The presence of spores was confirmed in seven out of ten of the larvae. At this time, fifty larvae were collected, washed with DI water, and homogenized in 1 mL of DI water using a glass pestle. The presence of binucleate spores in the homogenate was confirmed. Spores were then purified via Ludox density gradient assay as described in Solter et al., 2012. Briefly, the gradient was formed by layering 500 μL DI water on top of 500 μL Ludox reagent in a 1.5 ml microcentrifuge tube, laying the tube onto its side for 2 hours to permit gradient formation, then carefully returning the tube to an upright position to avoid disturbing the gradient. 300 μL of homogenate containing the spores was then layered on top of the gradient in the tube and centrifuged at 16,000 x g for 30 minutes. The resulting white band containing spores was harvested from the tube column under a flame and transferred to a 1.5 mL Eppendorf tube. This product was then washed (i.e., suspended in DI water, centrifuged at 1,500 x g for 10 minutes, and resuspended in fresh DI water after removal of supernatant) and resuspended in sterile DI water under a flame twice.

Following spore purification via Ludox density gradient assay, spore density was calculated using an inverted microscope and a hemocytometer and used for horizontal infections. The spore density of the final spore solution was 240,000 spores/mL. This spore solution was used to horizontally infect experimental larvae as described below. In addition, the spore solution was serially diluted ten-fold six times in sterile DI water. 100 μL of DNA was extracted from this dilution series using the ZymoBIOMICS™ DNA Miniprep Kit (Cat. #: D4300) following the manufacturer’s instructions and subjected to PCR using *E. aedis*-specific primers FQEA187 and RQEA310 (101) (Table S2). Each 25 μL reaction contained 10.5 μL molecular grade water, 0.5 μL of each primer (10 μM concentration), 1 μL of undiluted gDNA template, and 12.5 μL DreamTaq PCR master mix (Thermo Scientific™, Cat. #: FERK1081) and was performed on a T100 Thermal Cycler (Bio-Rad Cat. #: 1861096) under the following thermocycling conditions: 1) 95°C, 3 min, 2) 35 cycles of 95°C, 30 s; 55°C, 30 s; 72°C, 30 s, 3) 72°C, 10 min. The PCR product was run on a 1% agarose gel and the presence of a band of the correct size throughout the dilution series was used to identify a detection threshold by PCR of 240 spores (Fig. S2).

### Horizontal Edhazardia aedis infections & tissue collection

To query the effect of horizontal infection on the abdominal bacterial microbiota of non-blood-fed female *Ae. aegypti* (LVP strain), we used a paired mesocosm replication structure in which three paired sets of *E. aedis* (+)/control (mock-infected) larvae were reared in parallel from egg to adulthood (Fig. 1). Horizontal infection of experimental mosquito larvae with *E. aedis* was performed as described in (100). To prepare experimental mosquitoes, an unbleached egg paper was hatched in RO (reverse osmosis) water placed in a vacuum chamber. Upon hatching, larvae were thinned into three 12.75” x 10.44” x 2.5” polycarbonate trays (Cambro, Huntington Beach, CA) containing 1 L RO water at a density of 300 individuals per tray. At this time, 0.5 mL of tire water from an *Aedes* larval development site (tire water containing *Aedes* collected in Baltimore, MD September 2016 and stored in 25% glycerol at −70°C since collection) (102) was added to each tray for the purpose of inoculating the laboratory aquatic larval environment with a suite of microbes found in natural *Aedes* larval environments. An aliquot of this tire water inoculum was submitted for sequencing. At this time, each tray was given a pinch of fish food (Tetramin) and green cat food pellets (9 Lives) ad libitum. At 3 days post-hatching, larvae were randomly split and designated to an *E. aedis* infection group or a control (mock infection) group. To accomplish horizontal infection with *E. aedis*, 150 larvae were transferred to wide-mouth 500 mL clear borosilicate glass bottles (VWR Cat. #: 75871-304) containing 150 mL RO water and 3 mL freshly prepared liquid food slurry (liquid food slurry recipe: 1.2 g liver powder + 0.8 g brewer’s yeast powder + 100 mL DI water). Infection groups received 313 μL of the 240,000 spores/mL purified spore solution, which delivered approximately 75,000 spores to each bottle. Mock infection groups received the same treatment without the addition of *E. aedis* spores. Larvae were maintained in bottles for 24 hours. Following spore exposure in bottles, bottle contents were then transferred to 12.75” x 10.44” x 2.5” polycarbonate trays (Cambro, Huntington Beach, CA) and supplemented with 900mL RO water. Larvae were then reared with ad libitum access to green cat food pellets (9 Lives). Trays were checked daily to monitor pupation. Individuals that pupated between 6 and 8 days post-hatch were removed from trays daily with sterile transfer pipettes, washed with RO water, and transferred to sterile cups containing RO water and allowed to eclose. Adults were housed in 88 oz polypropylene container cages and provided with filter-sterilized 10% sucrose. The pupal water was replaced with fresh RO water daily until cups were removed.

Tissues were collected in a single instance when adults were aged 3-5 days post-eclosion. From each cage, n=7-8 individual females were aspirated and cold-anesthetized. Individuals were externally sterilized in a 3% bleach solution for 15 seconds, then rinsed twice with filter-sterilized 1X PBS. Abdomens were removed from the individuals beneath a microscope, placed into a 1.5mL microcentrifuge tube containing 150 μL lysis buffer from the ZymoBIOMICS™ DNA Miniprep Kit (Cat. #: D4300), homogenized, and stored at −80°C before DNA extraction per the manufacturer’s protocol. Dissection tools were cleaned with 70% ethanol and Kimwipes in between dissections.

We validated the infection status of *E. aedis* (+) females by subjecting all genomic mosquito DNA samples to PCR and gel electrophoresis using *E. aedis*-specific primers FQEA187 and RQEA310 (101) (Table S2) as described in the previous section. The assay confirmed the presence of *E. aedis* in all infected individuals (Fig. S3). The assay also confirmed the absence of infection in the control individuals; however, a faint signal is present in four out of the eight control individual samples in replicate 3 (Fig. S3). It is possible that DNA contamination or off-target DNA amplification led to this signal. However, given that the exclusion of these samples does not alter the findings, we do not believe that the source of this signal interfered with the validity of the dataset or biological findings presented.

### DNA extraction, library preparation, and sequencing data

Total DNA was extracted from mosquito abdomens, “blank” samples, and the tire water inoculum using the ZymoBIOMICS™ DNA Miniprep Kit (Cat. #: D4300) following the manufacturer’s instructions. Following DNA extraction, amplicon PCR was performed in-house using MyTaq™ Red Mix and primers 341F and 805R targeting the V3-V4 region of the 16S rRNA gene (103) (Table S2). Each 50 μL reaction contained 13 μL molecular grade water, 1 μL of each primer (10 μM concentration), 10 μL of undiluted gDNA template, and 25 μL MyTaq™ Red Mix PCR master mix (Meridian Bioscience, Cat. #: BIO-25043) and was performed on a T100 Thermal Cycler (Bio-Rad Cat. #: 1861096) under the following thermocycling conditions: 1) 95°C, 3 min, 2) 37 cycles of 95°C, 30 s; 55°C, 30 s; 72°C, 30 s, 3) 72°C, 10 min. PCR product was visualized on a 1% agarose gel and the target size amplicons were extracted using the Qiagen MinElute Gel Extraction kit (Cat. #: 28604) following the manufacturer’s instructions. Amplicons were submitted to NC State Genomic Sciences Laboratory for subsequent library preparation following the Illumina 16S Metagenomic Sequencing Library Preparation guide (104). Following library preparation, samples were sequenced on an Illumina NextSeq 2000 platform (300 base pair paired-ends reads). Reads were demultiplexed at NC State Genomic Sciences Laboratory with a 1 base pair error tolerance for each index.

### Bioinformatic analysis: quality control and data processing

Cutadapt (105) was used to trim primer and adapter contamination, as well as polyGs, from reads. Any forward reads containing reverse primers/adapters in the forward orientation or forward primers/adapters in the reverse complement orientation were then removed from the dataset using a custom code in R (File S1). The complementary process was repeated for the reverse reads. Phred quality score bins were identified as: 0, 9, 20, and 34. Following data cleanup, the DADA2 R package (106) was used for filtering, dereplication, sample inference, chimera identification, and merging of paired-end reads. Briefly: 1) Reads were first filtered with filterAndTrim() with default parameters without truncation, 2) Error rates were learned using default parameters, 3) The core dada2 sample inference algorithm was applied with pseudo-pooling, 4) reads were merged with a 20bp overlap requirement, 5) an amplicon sequence variant (ASV) table was constructed, 6) chimeras were removed, and 7) taxonomy was assigned using IDTAXA (107) trained against the SILVA SSU r138 training set (108) downloaded from DECIPHER Bioconductor package (109). A phyloseq object was created from the ASV table using the phyloseq package in R. ASVs with undefined kingdom-level classification and ASVs assigned to mitochondria or chloroplasts were removed from the dataset. Next, the decontam R package (110,111) was used to identify contaminants using the frequency method, which was implemented using index reaction DNA concentration values generated during sequencing library preparation. Finally, any ASVs that did not appear more than 10 times total in the dataset were removed from the phyloseq object. The resulting phyloseq object was used in all subsequent analyses. We recorded the number of reads lost at each step of quality control and data processing (Fig. S4, File S2). Code for all bioinformatic analysis can be found in File S1. The final phyloseq object used in subsequent analyses can be found in File S3.

### Statistical analysis of the sequencing dataset

All statistical analyses were performed using R version 4.4.1 (112) and R Studio version 2025.09.2 (113). R code for all statistical analyses can be found in File S1. The phyloseq object used in analyses can be found in File S3.

Ecological analyses were performed at multiple taxonomic levels. In all instances, merging of the dataset was accomplished using the tax_glom() function from the R package phyloseq (114).

To assess the effect(s) of *E. aedis* infection on alpha diversity of the microbiota, taxa richness was characterized using the Chao1 metric, while combined taxa richness and evenness was characterized using the Shannon and Simpson Indices. We then used Wilcoxon Rank-Sum tests to evaluate effects of infection on each alpha diversity index. We also used Kruskal-Wallis test to evaluate any variation in alpha diversity explained by larval mesocosm replicate.

To query the effect(s) of *E. aedis* infection on beta diversity, we constructed ordinations with four distances or transformed ASV tables: We constructed a PCoA using Bray-Curtis distances, as this is a common approach in the field. But because sequencing data is compositional data (CoDa) (73), we also constructed two compositional data-appropriate phylogeny-unaware ordinations with log-ratio transformed ASV tables and applied two different zero-handling approaches: 1) a robust centered log-ratio (rclr) transformation, in which only non-zero values are transformed, and 2) a centered log-ratio (clr) transformation on an ASV table in which the zeroes are imputed using Bayesian-multiplicative replacement, implemented using the cmultRepl() function from the zCompositions package (115). Finally, we used a CoDa-appropriate method to construct a phylogeny-aware ordination using balances derived from a phylogenetic isometric log-ratio (PhILR) transformation (116). We assessed the effect of *E. aedis* infection on the beta diversity of the microbiota using PERMANOVA tests, implemented via the adonis2() function from the R package vegan (117,118). We restricted permutations within each larval mesocosm replicate to account for any effects of replicate when assessing an overall effect of infection (119).

To query differential abundance of taxa, we used three differential abundance tools that are appropriate for compositional data: ANCOM-BC2 (120–122), ALDeX2 (123), and LinDA (124). All differential abundance analyses were performed at all taxonomic levels. In order for a taxon to be considered differentially abundant, we required identification that the taxon be identified by more than one tool.

We performed functional prediction analyses using FAPROTAX (125) and Tax4Fun2 (126).

### Bacterial load estimation using 16S qPCR

We also sought to understand the effect of *E. aedis* infection on bacterial load. Because sequencing data are compositional and cannot reflect the absolute abundance of starting material, we used quantitative PCR assays targeting the 16S rRNA bacterial gene and the *RpS7* mosquito gene (VectorBase: AAEL009496), with *RpS7* serving as the reference gene, to estimate the bacterial load in each sample. Reactions were performed on a Bio-Rad CFX96 Touch Deep Well Real-Time PCR Detection System in a volume of 10 μL containing 0.5 μL of each primer (the final concentration of each forward and reverse primer was 300nM for *RpS7* and 500nM for 16S), 4 μL of undiluted genomic DNA template, and 5 μL of PowerUp™ SYBR™ Green Master Mix for qPCR (Applied Biosystems™ Cat. #: A25741). The primer sequences were obtained from MacLeod et al., 2021 (102). Primer sequences can be found in Table S2. Cycling parameters for *RpS7* were: 2 min at 50°C + 2 min at 95°C + (15 sec at 95°C + 1 min at 56°C) × 40 cycles, followed by a dissociation (melt curve) step. Cycling parameters for 16S were: 2 min at 50°C + 2 min at 95°C + (15 sec at 95°C + 15 sec at 58°C + 1 min at 72°C) × 40 cycles, followed by a dissociation (melt curve) step. We performed melt curve analysis and agarose gel electrophoresis on representative qPCR products for both genes to validate the presence of a single product of the expected size. For each qPCR assay designed, we calculated an E_Amp_ (amplification efficiency) value using standard curves collected from at least three technical replicates. For *RpS7*, E_Amp,*RpS7*_ = 90%, and for 16S, E_Amp,16S_ = 95%. Standard curve dilution series were generated over a sufficiently wide range such that sample Cqs fell within the range of Cqs in the standard curves. Melt curve analysis was performed on all sample reactions and confirmed the presence of a single peak. Two or more technical replicates were performed per sample per gene with an inclusion criterion of 0.5 Cq distance from the average. We used the X_0_ method (127) to compute values reflecting the abundance of 16S DNA standardized to the abundance of *RpS7* DNA for each sample. Due to insufficient remaining genomic DNA template, we could not obtain qPCR data for 5 out of the 47 samples (three control samples and two *E. aedis*(+) samples), so these samples were excluded from analysis. Prior to analysis of ΔX_0,16S_, we confirmed that *E. aedis* treatment had no effect on the abundance of the reference gene *RpS7*. To assess the effect of *E. aedis* infection on bacterial load, we applied a log transformation to ΔX_0,16S_ values to improve normality, then built a mixed effects linear model using *E. aedis* infection status to predict log_10_(ΔX_0,16S_) with random intercepts for larval mesocosm replicates. The qPCR data can be found in File S4.

## Data availability statement

Sequence data have been submitted to the Sequence Read Archive under accession number PRJNA1415287.

## Supporting information legends

**Fig. S1. Rarefaction curves generated from the final phyloseq object**.

**Fig. S2. Establishing *E. aedis* limit of detection with PCR and gel electrophoresis**. 100 μL DNA was extracted from each sample in a tenfold dilution series of the final spore solution (240,000 spores/mL) used for horizontal *E. aedis* infections and subjected to PCR with *E. aedis*-specific primers FQEA187/RQEA310 and gel electrophoresis analysis to establish a PCR detection threshold of 24 spores. 1 Kb Plus DNA Ladder was used (Invitrogen Cat. #: 10787018).

**Fig. S3. *E. aedis* infection confirmation in mosquito samples with PCR and gelelectrophoresis**. Mosquito abdomen genomic DNA was subjected to PCR with *E. aedis*-specific primers FQEA187/RQEA310 and gel electrophoresis to confirm infection status of mosquitoes. The infection status of sample “Mock1.2” was not assessed due to insufficient remaining sample volume. 1 Kb Plus DNA Ladder was used (Invitrogen Cat. #: 10787018).

**Fig. S4. Displaying the number of reads lost at each step of bioinformatic data processing**. The data are stored in a .csv file (File S2) and the R code used to generate the figure is in File S1.

**File S1. R code used for all data processing, analysis, and visualization**.

**File S2. A .csv file storing the number of reads lost at each step of bioinformatic data processing**. The data were used to generate Fig. S4 using R code in File S1.

**File S3. The final phyloseq object used in analyses. File S4. A .csv file storing the qPCR data**.

**Table S1. Taxonomic information for “Core” ASVs (present in all n=47 mosquito samples)**. Asterisks denote ASVs that were also identified in tire water inoculum.

**Table S2. Primer Information**. An asterisk indicates the presence of sequencing overhang adapters in the sequences, which are denoted in bold.

## Acknowledgements

We wish to thank Dr. James Becnel and Dr. Neil Sanscrainte for providing *E. aedis* spores.

## References

1. Greer R, Dong X, Morgun A, Shulzhenko N. Investigating a holobiont: Microbiota perturbations and transkingdom networks. Gut Microbes. 2016 Mar 3;7(2):126–35.

2. Simon JC, Marchesi JR, Mougel C, Selosse MA. Host-microbiota interactions: from holobiont theory to analysis. Microbiome. 2019 Dec;7(1):5.

3. Belkaid Y, Harrison OJ. Homeostatic Immunity and the Microbiota. Immunity. 2017 Apr;46(4):562–76.

4. Debnath N, Kumar R, Kumar A, Mehta PK, Yadav AK. Gut-microbiota derived bioactive metabolites and their functions in host physiology. Biotechnol Genet Eng Rev. 2021 Jul 3;37(2):105–53.

5. Ikeda-Ohtsubo W, Brugman S, Warden CH, Rebel JMJ, Folkerts G, Pieterse CMJ. How Can We Define “Optimal Microbiota?”: A Comparative Review of Structure and Functions of Microbiota of Animals, Fish, and Plants in Agriculture. Front Nutr. 2018 Oct 2;5:90.

6. Petersen C, Round JL. Defining dysbiosis and its influence on host immunity and disease. Cell Microbiol. 2014 Jul;16(7):1024–33.

7. Ram H, Dastager SG. Re-purposing is needed for beneficial bugs, not for the drugs. Int Microbiol. 2019 Mar;22(1):1–6.

8. Rosenfeld CS. Gut Dysbiosis in Animals Due to Environmental Chemical Exposures. Front Cell Infect Microbiol. 2017 Sep 8;7:396.

9. Flores GAM, Lopez RP, Cerrudo CS, Consolo VF, Berón CM. Culex quinquefasciatus Holobiont: A Fungal Metagenomic Approach. Front Fungal Biol. 2022 Aug 2;3:918052.

10. Guégan M, Zouache K, Démichel C, Minard G, Tran Van V, Potier P, et al. The mosquito holobiont: fresh insight into mosquito-microbiota interactions. Microbiome. 2018 Dec;6(1):49.

11. Minard G, Mavingui P, Moro CV. Diversity and function of bacterial microbiota in the mosquito holobiont. Parasit Vectors. 2013 Dec;6(1):146.

12. Singh U, Das A. Vector-borne diseases: Mosquito holobiont and novel methods for vector control. Asian Pac J Trop Biomed. 2020;10(8):341.

13. Zheng R, Wang Q, Wu R, Paradkar PN, Hoffmann AA, Wang GH. Holobiont perspectives on tripartite interactions among microbiota, mosquitoes, and pathogens. ISME J. 2023 Aug 1;17(8):1143–52.

14. Crawford JE, Clarke DW, Criswell V, Desnoyer M, Cornel D, Deegan B, et al. Efficient production of male Wolbachia-infected Aedes aegypti mosquitoes enables large-scale suppression of wild populations. Nat Biotechnol. 2020 Apr 1;38(4):482–92.

15. Patyka TI, Patyka, M.V. Bacillus thuringiensis spp. israelensis and Control of Aedes aegypti Invasive Mosquitoes Species in Ecosystems. Mikrobiol Zh. 2020 Oct 17;82(5):88–97.

16. The Lancet. Dengue: the threat to health now and in the future. The Lancet. 2024 Jul;404(10450):311.

17. Wang Y, Gilbreath TM, Kukutla P, Yan G, Xu J. Dynamic Gut Microbiome across Life History of the Malaria Mosquito Anopheles gambiae in Kenya. Leulier F, editor. PLoS ONE. 2011 Sep 21;6(9):e24767.

18. Coon KL, Vogel KJ, Brown MR, Strand MR. Mosquitoes rely on their gut microbiota for development. Mol Ecol. 2014 Jun;23(11):2727–39.

19. Arévalo-Cortés A, Mejia-Jaramillo AM, Granada Y, Coatsworth H, Lowenberger C, Triana-Chavez O. The Midgut Microbiota of Colombian Aedes aegypti Populations with Different Levels of Resistance to the Insecticide Lambda-cyhalothrin. Insects. 2020 Sep 1;11(9):584.

20. Dada N, Sheth M, Liebman K, Pinto J, Lenhart A. Whole metagenome sequencing reveals links between mosquito microbiota and insecticide resistance in malaria vectors. Sci Rep. 2018 Feb 1;8(1):2084.

21. Omoke D, Kipsum M, Otieno S, Esalimba E, Sheth M, Lenhart A, et al. Western Kenyan Anopheles gambiae showing intense permethrin resistance harbour distinct microbiota. Malar J. 2021 Feb 8;20(1):77.

22. Pelloquin B, Kristan M, Edi C, Meiwald A, Clark E, Jeffries CL, et al. Overabundance of Asaia and Serratia Bacteria Is Associated with Deltamethrin Insecticide Susceptibility in Anopheles coluzzii from Agboville, Côte d’Ivoire. Garg N, editor. Microbiol Spectr. 2021 Oct 31;9(2):e00157–21.

23. Soltani A, Vatandoost H, Oshaghi MA, Enayati AA, Chavshin AR. The role of midgut symbiotic bacteria in resistance of Anopheles stephensi (Diptera: Culicidae) to organophosphate insecticides. Pathog Glob Health. 2017 Aug 18;111(6):289–96.

24. Rodgers FH, Gendrin M, Wyer CAS, Christophides GK. Microbiota-induced peritrophic matrix regulates midgut homeostasis and prevents systemic infection of malaria vector mosquitoes. Dimopoulos G, editor. PLOS Pathog. 2017 May 17;13(5):e1006391.

25. Gaio ADO, Gusmão DS, Santos AV, Berbert-Molina MA, Pimenta PF, Lemos FJ. Contribution of midgut bacteria to blood digestion and egg production in Aedes aegypti (diptera: culicidae) (L.). Parasit Vectors. 2011 Dec;4(1):105.

26. Guégan M, Tran Van V, Martin E, Minard G, Tran F, Fel B, et al. Who is eating fructose within the A<SC>EDES ALBOPICTUS </SC>gut microbiota? Environ Microbiol. 2020 Apr;22(4):1193–206.

27. Guégan M, Martin E, Tran Van V, Fel B, Hay AE, Simon L, et al. Mosquito sex and mycobiota contribute to fructose metabolism in the Asian tiger mosquito Aedes albopictus. Microbiome. 2022 Aug 30;10(1):138.

28. Salgado JFM, Premkrishnan BNV, Oliveira EL, Vettath VK, Goh FG, Hou X, et al. The dynamics of the midgut microbiome in Aedes aegypti during digestion reveal putative symbionts. Dupont C, editor. PNAS Nexus. 2024 Aug 5;3(8):pgae317.

29. Villegas LEM, Radl J, Dimopoulos G, Short SM. Bacterial communities of Aedes aegypti mosquitoes differ between crop and midgut tissues. Fischer K, editor. PLoS Negl Trop Dis. 2023 Mar 29;17(3):e0011218.

30. Mosquera KD, Martínez Villegas LE, Rocha Fernandes G, Rocha David M, Maciel-de-Freitas R, A Moreira L, et al. Egg-laying by female Aedes aegypti shapes the bacterial communities of breeding sites. BMC Biol. 2023 Apr 26;21(1):97.

31. Angleró-Rodríguez YI, Blumberg BJ, Dong Y, Sandiford SL, Pike A, Clayton AM, et al. A natural Anopheles-associated Penicillium chrysogenum enhances mosquito susceptibility to Plasmodium infection. Sci Rep. 2016 Sep 28;6(1):34084.

32. Apte-Deshpande AD, Paingankar MS, Gokhale MD, Deobagkar DN. Serratia odorifera mediated enhancement in susceptibility of Aedes aegypti for chikungunya virus. Indian J Med Res. 2014 May;139(5):762.

33. Bolling BG, Olea-Popelka FJ, Eisen L, Moore CG, Blair CD. Transmission dynamics of an insect-specific flavivirus in a naturally infected Culex pipiens laboratory colony and effects of co-infection on vector competence for West Nile virus. Virology. 2012 Jun;427(2):90–7.

34. Bongio NJ, Lampe DJ. Inhibition of Plasmodium berghei Development in Mosquitoes by Effector Proteins Secreted from Asaia sp. Bacteria Using a Novel Native Secretion Signal. Favia G, editor. PLOS ONE. 2015 Dec 4;10(12):e0143541.

35. Cirimotich CM, Dong Y, Clayton AM, Sandiford SL, Souza-Neto JA, Mulenga M, et al. Natural Microbe-Mediated Refractoriness to Plasmodium Infection in Anopheles gambiae. Science. 2011 May 13;332(6031):855–8.

36. Dickson LB, Jiolle D, Minard G, Moltini-Conclois I, Volant S, Ghozlane A, et al. Carryover effects of larval exposure to different environmental bacteria drive adult trait variation in a mosquito vector. Sci Adv. 2017 Aug 4;3(8):e1700585.

37. Goenaga S, Kenney J, Duggal N, Delorey M, Ebel G, Zhang B, et al. Potential for Co-Infection of a Mosquito-Specific Flavivirus, Nhumirim Virus, to Block West Nile Virus Transmission in Mosquitoes. Viruses. 2015 Nov 11;7(11):5801–12.

38. Herren JK, Mbaisi L, Mararo E, Makhulu EE, Mobegi VA, Butungi H, et al. A microsporidian impairs Plasmodium falciparum transmission in Anopheles arabiensis mosquitoes. Nat Commun. 2020 May 4;11(1):2187.

39. Ramirez JL, Short SM, Bahia AC, Saraiva RG, Dong Y, Kang S, et al. Chromobacterium Csp_P Reduces Malaria and Dengue Infection in Vector Mosquitoes and Has Entomopathogenic and In Vitro Anti-pathogen Activities. Levashina E, editor. PLoS Pathog. 2014 Oct 23;10(10):e1004398.

40. Tchioffo MT, Boissière A, Churcher TS, Abate L, Gimonneau G, Nsango SE, et al. Modulation of Malaria Infection in Anopheles gambiae Mosquitoes Exposed to Natural Midgut Bacteria. Michel K, editor. PLoS ONE. 2013 Dec 6;8(12):e81663.

41. Valzano M, Cecarini V, Cappelli A, Capone A, Bozic J, Cuccioloni M, et al. A yeast strain associated to Anopheles mosquitoes produces a toxin able to kill malaria parasites. Malar J. 2016 Dec;15(1):21.

42. Wang Y, Wang Y, Zhang J, Xu W, Zhang J, Huang FS. Ability of TEP1 in intestinal flora to modulate natural resistance of Anopheles dirus. Exp Parasitol. 2013 Aug;134(4):460–5.

43. Ruan Y, Xu X, He Q, Li L, Guo J, Bao J, et al. The largest meta-analysis on the global prevalence of microsporidia in mammals, avian and water provides insights into the epidemic features of these ubiquitous pathogens. Parasit Vectors. 2021 Apr 1;14(1):186.

44. Becnel JJ, Andreadis TG. Microsporidia in Insects. In: Weiss LM, Becnel JJ, editors. Microsporidia [Internet]. 1st ed. Wiley; 2014 [cited 2024 Oct 2]. p. 521–70. Available from: https://onlinelibrary.wiley.com/doi/10.1002/9781118395264.ch21

45. Hembree, Stephen C. Preliminary report of some mosquito pathogens from Thailand. Mosquito News. 1979 Sep;39(3):575–82.

46. Kudo, Richard Roksabro. Studies on Microsporidia parasitic in mosquitoes. VIII. On a microsporidian, Nosema aedis nov. spec., parasitic in a larva of Aedes aegypti of Porto Rico. Archiv für Protistenkunde. 1930;69.

47. Nasci RS, Tang KH, Becnel JJ, Fukuda T. Effect of per os Edhazardia aedis (Microsporida: Amblyosporidae) infection on Aedes aegypti mortality and body size. J Am Mosq Control Assoc. 1992 Jun;8(2):131–6.

48. Becnel JJ, Sprague V, Fukuda T, Hazard EI. Development of Edhazardia aedis (Kudo, 1930) N. G., N. Comb. (Microsporida: Amblyosporidae) in the Mosquito Aedes aegypti (L.) (Diptera: Culicidae). J Protozool. 1989 Mar;36(2):119–30.

49. Agnew P, Koella JC. Life history interactions with environmental conditions in a host–parasite relationship and the parasite’s mode of transmission. Evol Ecol. 1999 Jan;13(1):67–91.

50. Zilio G, Kaltz O, Koella JC. Resource availability for the mosquito Aedes aegypti affects the transmission mode evolution of a microsporidian parasite. Evol Ecol. 2023 Feb;37(1):31–51.

51. Desjardins CA, Sanscrainte ND, Goldberg JM, Heiman D, Young S, Zeng Q, et al. Contrasting host–pathogen interactions and genome evolution in two generalist and specialist microsporidian pathogens of mosquitoes. Nat Commun. 2015 May 13;6(1):7121.

52. El-Dougdoug NK, Magistrado D, Short SM. An obligate microsporidian parasite modulates defense against opportunistic bacterial infection in the yellow fever mosquito, Aedes aegypti. Blader IJ, editor. mSphere. 2024 Feb 28;9(2):e00678–23.

53. Barnard DR, Xue RD, Rotstein MA, Becnel JJ. Microsporidiosis (Microsporidia: Culicosporidae) Alters Blood-Feeding Responses and DEET Repellency in Aedes aegypti (Diptera: Culicidae). J Med Entomol. 2007 Nov 1;44(6):1040–6.

54. Becnel JJ, Garcia JJ, Johnson MA. Edhazardia aedis (Microspora: Culicosporidae) Effects on the Reproductive Capacity of Aedes aegypti (Diptera: Culicidae). J Med Entomol. 1995 Jul 1;32(4):549–53.

55. Koella JC, Agnew P. Blood-Feeding Success of the Mosquito Aedes aegypti Depends on the Transmission Route of Its Parasite Edhazardia aedis. Oikos. 1997 Mar;78(2):311.

56. Andreadis TG. Host Range Tests with Edhazardia aedis (Microsporida: Culicosporidae) against Northern Nearctic Mosquitoes. J Invertebr Pathol. 1994 Jul;64(1):46–51.

57. Becnel JJ, Johnson MA. Mosquito host range and specificity of Edhazardia aedis (Microspora: Culicosporidae). J Am Mosq Control Assoc. 1993 Sep;9(3):269–74.

58. Brimner TA, Boland GJ. A review of the non-target effects of fungi used to biologically control plant diseases. Agric Ecosyst Environ. 2003 Nov;100(1):3–16.

59. Perazzolli M, Antonielli L, Storari M, Puopolo G, Pancher M, Giovannini O, et al. Resilience of the Natural Phyllosphere Microbiota of the Grapevine to Chemical and Biological Pesticides. Drake HL, editor. Appl Environ Microbiol. 2014 Jun 15;80(12):3585–96.

60. Scheepmaker JWA, Kassteele JVD. Effects of chemical control agents and microbial biocontrol agents on numbers of non-target microbial soil organisms: a meta-analysis. Biocontrol Sci Technol. 2011 Oct;21(10):1225–42.

61. Alberoni D, Di Gioia D, Baffoni L. Alterations in the Microbiota of Caged Honeybees in the Presence of Nosema ceranae Infection and Related Changes in Functionality. Microb Ecol. 2023 Jul;86(1):601–16.

62. Boissière A, Tchioffo MT, Bachar D, Abate L, Marie A, Nsango SE, et al. Midgut Microbiota of the Malaria Mosquito Vector Anopheles gambiae and Interactions with Plasmodium falciparum Infection. Vernick KD, editor. PLoS Pathog. 2012 May 31;8(5):e1002742.

63. Maes PW, Rodrigues PAP, Oliver R, Mott BM, Anderson KE. Diet-related gut bacterial dysbiosis correlates with impaired development, increased mortality and Nosema disease in the honeybee (Apis mellifera). Mol Ecol. 2016 Nov;25(21):5439–50.

64. Mousson L, Martin E, Zouache K, Madec Y, Mavingui P, Failloux AB. Wolbachia modulates Chikungunya replication in Aedes albopictus. Mol Ecol. 2010 May;19(9):1953–64.

65. Naree S, Ellis JD, Benbow ME, Suwannapong G. Experimental Nosema ceranae infection is associated with microbiome changes in the midguts of four species of Apis (honey bees). J Apic Res. 2022 May 27;61(3):435–47.

66. Panjad P, Yongsawas R, Sinpoo C, Pakwan C, Subta P, Krongdang S, et al. Impact of Nosema Disease and American Foulbrood on Gut Bacterial Communities of Honeybees Apis mellifera. Insects. 2021 Jun 6;12(6):525.

67. Paris L, Peghaire E, Moné A, Diogon M, Debroas D, Delbac F, et al. Honeybee gut microbiota dysbiosis in pesticide/parasite co-exposures is mainly induced by Nosema ceranae. J Invertebr Pathol. 2020 May;172:107348.

68. Shen H, Dou Y, Li H, Qiao Y, Jiang G, Wan X, et al. Changes in the intestinal microbiota of Pacific white shrimp (Litopenaeus vannamei) with different severities of Enterocytozoon hepatopenaei infection. J Invertebr Pathol. 2022 Jun;191:107763.

69. Villegas LEM, Campolina TB, Barnabe NR, Orfano AS, Chaves BA, Norris DE, et al. Zika virus infection modulates the bacterial diversity associated with Aedes aegypti as revealed by metagenomic analysis. Oliveira PL, editor. PLOS ONE. 2018 Jan 2;13(1):e0190352.

70. Wei G, Lai Y, Wang G, Chen H, Li F, Wang S. Insect pathogenic fungus interacts with the gut microbiota to accelerate mosquito mortality. Proc Natl Acad Sci. 2017 Jun 6;114(23):5994–9.

71. Zhang X, Feng H, He J, Liang X, Zhang N, Shao Y, et al. The gut commensal bacterium E<SC>NTEROCOCCUS FAECALIS </SC>LX10 contributes to defending against N<SC>OSEMA BOMBYCIS </SC>infection in B<SC>OMBYX MORI </SC>. Pest Manag Sci. 2022 Jun;78(6):2215–27.

72. Zouache K, Michelland RJ, Failloux A, Grundmann GL, Mavingui P. Chikungunya virus impacts the diversity of symbiotic bacteria in mosquito vector. Mol Ecol. 2012 May;21(9):2297–309.

73. Gloor GB, Macklaim JM, Pawlowsky-Glahn V, Egozcue JJ. Microbiome Datasets Are Compositional: And This Is Not Optional. Front Microbiol. 2017 Nov 15;8:2224.

74. Boissière A, Tchioffo MT, Bachar D, Abate L, Marie A, Nsango SE, et al. Midgut Microbiota of the Malaria Mosquito Vector Anopheles gambiae and Interactions with Plasmodium falciparum Infection. Vernick KD, editor. PLoS Pathog. 2012 May 31;8(5):e1002742.

75. Mousson L, Martin E, Zouache K, Madec Y, Mavingui P, Failloux AB. Wolbachia modulates Chikungunya replication in Aedes albopictus. Mol Ecol. 2010 May;19(9):1953–64.

76. Wei G, Lai Y, Wang G, Chen H, Li F, Wang S. Insect pathogenic fungus interacts with the gut microbiota to accelerate mosquito mortality. Proc Natl Acad Sci. 2017 Jun 6;114(23):5994–9.

77. Kamada N, Seo SU, Chen GY, Núñez G. Role of the gut microbiota in immunity and inflammatory disease. Nat Rev Immunol. 2013 May;13(5):321–35.

78. Hixson B, Chen R, Buchon N. Innate immunity in Aedes mosquitoes: from pathogen resistance to shaping the microbiota. Philos Trans R Soc B Biol Sci. 2024 May 6;379(1901):20230063.

79. Byndloss MX, Pernitzsch SR, Bäumler AJ. Healthy hosts rule within: ecological forces shaping the gut microbiota. Mucosal Immunol. 2018 Sep;11(5):1299–305.

80. Litvak Y, Byndloss MX, Bäumler AJ. Colonocyte metabolism shapes the gut microbiota. Science. 2018 Nov 30;362(6418):eaat9076.

81. McLoughlin K, Schluter J, Rakoff-Nahoum S, Smith AL, Foster KR. Host Selection of Microbiota via Differential Adhesion. Cell Host Microbe. 2016 Apr;19(4):550–9.

82. Schluter J, Foster KR. The Evolution of Mutualism in Gut Microbiota Via Host Epithelial Selection. Ellner SP, editor. PLoS Biol. 2012 Nov 20;10(11):e1001424.

83. Wilde J, Slack E, Foster KR. Host control of the microbiome: Mechanisms, evolution, and disease. Science. 2024 Jul 19;385(6706):eadi3338.

84. Kosakamoto H, Yamauchi T, Akuzawa-Tokita Y, Nishimura K, Soga T, Murakami T, et al. Local Necrotic Cells Trigger Systemic Immune Activation via Gut Microbiome Dysbiosis in Drosophila. Cell Rep. 2020 Jul;32(3):107938.

85. Negroni MA, Segers FHID, Vogelweith F, Foitzik S. Immune challenge reduces gut microbial diversity and triggers fertility-dependent gene expression changes in a social insect. BMC Genomics. 2020 Dec;21(1):816.

86. Buttó LF, Haller D. Dysbiosis in intestinal inflammation: Cause or consequence. Int J Med Microbiol. 2016 Aug;306(5):302–9.

87. Winter SE, Bäumler AJ. Gut dysbiosis: Ecological causes and causative effects on human disease. Proc Natl Acad Sci. 2023 Dec 12;120(50):e2316579120.

88. Dey P, Ray Chaudhuri S. The opportunistic nature of gut commensal microbiota. Crit Rev Microbiol. 2023 Nov 2;49(6):739–63.

89. Trzebny A, Slodkowicz-Kowalska A, Björkroth J, Dabert M. Microsporidian Infection in Mosquitoes (Culicidae) Is Associated with Gut Microbiome Composition and Predicted Gut Microbiome Functional Content. Microb Ecol. 2023 Jan;85(1):247–63.

90. Onchuru TO, Makhulu EE, Ronnie PC, Mandere S, Otieno FG, Gichuhi J, et al. The Plasmodium transmission-blocking symbiont, Microsporidia MB, is vertically transmitted through Anopheles arabiensis germline stem cells. Ferreira MU, editor. PLOS Pathog. 2024 Nov 11;20(11):e1012340.

91. Foster WA. Mosquito Sugar Feeding and Reproductive Energetics. Annu Rev Entomol. 1995 Jan;40(1):443–74.

92. Hurd H, Hogg JC, Renshaw M. Interactions between bloodfeeding, fecundity and infection in mosquitoes. Parasitol Today. 1995 Nov;11(11):411–6.

93. Kelly BJ, Joseph NC, Magistrado D, Short SM. The effects of mating and blood feeding on the immune defense of female Aedes aegypti mosquitoes. Abd-Alla AMM, editor. PLoS Negl Trop Dis. 2025 Oct 3;19(10):e0013542.

94. Magistrado D, El-Dougdoug NK, Short SM. Sugar restriction and blood ingestion shape divergent immune defense trajectories in the mosquito Aedes aegypti. Sci Rep. 2023 Jul 31;13(1):12368.

95. Santiago PB, D. Araújo CN, Motta FN, Praça YR, Charneau S, Bastos IMD, et al. Proteases of haematophagous arthropod vectors are involved in blood-feeding, yolk formation and immunity - a review. Parasit Vectors. 2017 Dec;10(1):79.

96. Schwenke RA, Lazzaro BP, Wolfner MF. Reproduction–Immunity Trade-Offs in Insects. Annu Rev Entomol. 2016 Mar 11;61(1):239–56.

97. Gusmão DS, Santos AV, Marini DC, Bacci M, Berbert-Molina MA, Lemos FJA. Culture-dependent and culture-independent characterization of microorganisms associated with Aedes aegypti (Diptera: Culicidae) (L.) and dynamics of bacterial colonization in the midgut. Acta Trop. 2010 Sep;115(3):275–81.

98. Muturi EJ, Njoroge TM, Dunlap C, Cáceres CE. Blood meal source and mixed blood-feeding influence gut bacterial community composition in Aedes aegypti. Parasit Vectors. 2021 Dec;14(1):83.

99. Oliveira JHM, Gonçalves RLS, Lara FA, Dias FA, Gandara ACP, Menna-Barreto RFS, et al. Blood Meal-Derived Heme Decreases ROS Levels in the Midgut of Aedes aegypti and Allows Proliferation of Intestinal Microbiota. Schneider DS, editor. PLoS Pathog. 2011 Mar 17;7(3):e1001320.

100. Grigsby A, Kelly BJ, Sanscrainte ND, Becnel JJ, Short SM. Propagation of the Microsporidian Parasite Edhazardia aedis in Aedes aegypti Mosquitoes. J Vis Exp. 2020 Aug 13;(162):61574.

101. Duncan AB, Agnew P, Noel V, Demettre E, Seveno M, Brizard J, et al. Proteome of Aedes aegypti in response to infection and coinfection with microsporidian parasites. Ecol Evol. 2012 Apr;2(4):681–94.

102. MacLeod HJ, Dimopoulos G, Short SM. Larval Diet Abundance Influences Size and Composition of the Midgut Microbiota of Aedes aegypti Mosquitoes. Front Microbiol. 2021 Jun 18;12:645362.

103. Klindworth A, Pruesse E, Schweer T, Peplies J, Quast C, Horn M, et al. Evaluation of general 16S ribosomal RNA gene PCR primers for classical and next-generation sequencing-based diversity studies. Nucleic Acids Res. 2013 Jan 1;41(1):e1–e1.

104. 16S Metagenomic Sequencing Library Preparation: Preparing 16S Ribosomal RNA Gene Amplicons for the Illumina MiSeq System. Illumina; 2013 Nov. Report No.: Part # 15044223 Rev. B.

105. Martin M. Cutadapt removes adapter sequences from high-throughput sequencing reads. EMBnet.journal. 2011 May 2;17(1):10.

106. Callahan BJ, McMurdie PJ, Rosen MJ, Han AW, Johnson AJA, Holmes SP. DADA2: High-resolution sample inference from Illumina amplicon data. Nat Methods. 2016 Jul;13(7):581–3.

107. Murali A, Bhargava A, Wright ES. IDTAXA: a novel approach for accurate taxonomic classification of microbiome sequences. Microbiome. 2018 Dec;6(1):140.

108. Yilmaz P, Parfrey LW, Yarza P, Gerken J, Pruesse E, Quast C, et al. The SILVA and “All-species Living Tree Project (LTP)” taxonomic frameworks. Nucleic Acids Res. 2014 Jan;42(D1):D643–8.

109. Wright E. DECIPHER [Internet]. Bioconductor; 2017 [cited 2025 Jan 23]. Available from: https://bioconductor.org/packages/DECIPHER

110. Benjamin Callahan NMD. decontam [Internet]. Bioconductor; 2018 [cited 2024 Nov 19]. Available from: https://bioconductor.org/packages/decontam

111. Davis NM, Proctor DM, Holmes SP, Relman DA, Callahan BJ. Simple statistical identification and removal of contaminant sequences in marker-gene and metagenomics data [Internet]. Bioinformatics; 2017 [cited 2024 Nov 19]. Available from: http://biorxiv.org/lookup/doi/10.1101/221499

112. R Core Team. R: A language and environment for statistical computing [Internet]. Vienna, Austria: R Foundation for Statistical Computing; 2025. Available from: https://www.R-project.org/

113. Posit team. RStudio: Integrated Development Environment for R [Internet]. Boston, MA: Posit Software; 2025. Available from: http://www.posit.co/

114. McMurdie PJ, Holmes S. phyloseq: An R Package for Reproducible Interactive Analysis and Graphics of Microbiome Census Data. Watson M, editor. PLoS ONE. 2013 Apr 22;8(4):e61217.

115. Palarea-Albaladejo J, Martín-Fernández JA. zCompositions — R package for multivariate imputation of left-censored data under a compositional approach. Chemom Intell Lab Syst. 2015 Apr;143:85–96.

116. Silverman JD, Washburne AD, Mukherjee S, David LA. A phylogenetic transform enhances analysis of compositional microbiota data. eLife. 2017 Feb 15;6:e21887.

117. Dixon P. VEGAN, a package of R functions for community ecology. J Veg Sci. 2003 Dec;14(6):927–30.

118. Oksanen J, Simpson GL, Blanchet FG, Kindt R, Legendre P, Minchin PR, et al. vegan: Community Ecology Package [Internet]. 2001 [cited 2025 Dec 21]. p. 2. 7–2. Available from: https://CRAN.R-project.org/package=vegan

119. Anderson M, Braak CT. Permutation tests for multi-factorial analysis of variance. J Stat Comput Simul. 2003 Jan;73(2):85–113.

120. Lin H, Eggesbø M, Peddada SD. Linear and nonlinear correlation estimators unveil undescribed taxa interactions in microbiome data. Nat Commun. 2022 Aug 23;13(1):4946.

121. Lin H, Peddada SD. Analysis of compositions of microbiomes with bias correction. Nat Commun. 2020 Jul 14;11(1):3514.

122. Lin H, Peddada SD. Multigroup analysis of compositions of microbiomes with covariate adjustments and repeated measures. Nat Methods. 2024 Jan;21(1):83–91.

123. Gloor GB, Macklaim JM, Fernandes AD. Displaying Variation in Large Datasets: Plotting a Visual Summary of Effect Sizes. J Comput Graph Stat. 2016 Jul 2;25(3):971–9.

124. Zhou H, He K, Chen J, Zhang X. LinDA: linear models for differential abundance analysis of microbiome compositional data. Genome Biol. 2022 Dec;23(1):95.

125. Louca S, Parfrey LW, Doebeli M. Decoupling function and taxonomy in the global ocean microbiome. Science. 2016 Sep 16;353(6305):1272–7.

126. Wemheuer F, Taylor JA, Daniel R, Johnston E, Meinicke P, Thomas T, et al. Tax4Fun2: prediction of habitat-specific functional profiles and functional redundancy based on 16S rRNA gene sequences. Environ Microbiome. 2020 May 18;15(1):11.

127. Thomsen R, Sølvsten CAE, Linnet TE, Blechingberg J, Nielsen AL. Analysis Of Qpcr Data By Converting Exponentially Related Ct Values Into Linearly Related X0 Values. J Bioinform Comput Biol. 2010 Oct;08(05):885–900.

