## Supplementary material for "The *Aedes aegypti* bacterial microbiota is robust to infection with the obligate microsporidian parasite *Edhazardia aedis*": Table S1

**Table S1. Taxonomic information for “Core” ASVs (present in all n=47 mosquito samples).** Asterisks denote ASVs that were also identified in tire water inoculum.

| **ASV** | **Phylum** | **Class** | **Order** | **Family** | **Genus** |
| --- | --- | --- | --- | --- | --- |
| ASV1* | Pseudomonadota | Gammaproteobacteria | Pseudomonadales | Pseudomonadaceae | Pseudomonas |
| ASV2* | Bacillota | Bacilli | Staphylococcales | Staphylococcaceae | Staphylococcus |
| ASV3* | Pseudomonadota | Gammaproteobacteria | Burkholderiales | Comamonadaceae | NA |
| ASV4* | Pseudomonadota | Gammaproteobacteria | Burkholderiales | Burkholderiaceae | Burkholderia-Caballeronia-Paraburkholderia |
| ASV6* | Pseudomonadota | Gammaproteobacteria | Pseudomonadales | Moraxellaceae | Acinetobacter |
| ASV7* | Bacillota | Bacilli | Staphylococcales | Staphylococcaceae | Staphylococcus |
| ASV8* | Pseudomonadota | Alphaproteobacteria | Sphingomonadales | Sphingomonadaceae | NA |
| ASV9* | Bacillota | Bacilli | Staphylococcales | Staphylococcaceae | Staphylococcus |
| ASV10 | Bacillota | Bacilli | Staphylococcales | Staphylococcaceae | Staphylococcus |
| ASV12 | Pseudomonadota | Gammaproteobacteria | Burkholderiales | Burkholderiaceae | Burkholderia-Caballeronia-Paraburkholderia |
| ASV14* | Pseudomonadota | Alphaproteobacteria | Azospirillales | Azospirillaceae | Nitrospirillum |
| ASV15* | Bacillota | Bacilli | Bacillales | Planococcaceae | Solibacillus |
| ASV16* | Pseudomonadota | Gammaproteobacteria | Pseudomonadales | Moraxellaceae | Psychrobacter |
| ASV19* | Pseudomonadota | Alphaproteobacteria | Acetobacterales | Acetobacteraceae | Asaia |
| ASV21* | Pseudomonadota | Gammaproteobacteria | Burkholderiales | Comamonadaceae | NA |
| ASV26 | Pseudomonadota | Gammaproteobacteria | Burkholderiales | Oxalobacteraceae | Herbaspirillum |
| ASV31 | Bacillota | Bacilli | Lactobacillales | Carnobacteriaceae | Desemzia |
| ASV33* | Pseudomonadota | Gammaproteobacteria | Pseudomonadales | Moraxellaceae | Acinetobacter |
| ASV35* | Actinomycetota | Actinobacteria | Propionibacteriales | Propionibacteriaceae | Cutibacterium |
| ASV321* | Pseudomonadota | Gammaproteobacteria | Enterobacterales | Yersiniaceae | NA |
