## Supplementary material for "The *Aedes aegypti* bacterial microbiota is robust to infection with the obligate microsporidian parasite *Edhazardia aedis*": Table S2

**Table S2. Primer Information.** An asterisk indicates the presence of sequencing overhang adapters in the sequences, which are denoted in bold.

| Assay | Primer name | Sequence (5’ - 3’) | Source (see Bibliography) |
| --- | --- | --- | --- |
| PCR amplicon reaction | 341F* | **TCGTCGGCAGCGTCAGATGTGTATAAGAGACAG**CCTACGGGNGGCWGCAG | Klindworth et al., 2012  (<doi.org/10.1093/nar/gks808>) |
|  | 805R* | **GTCTCGTGGGCTCGGAGATGTGTATAAGAGACAG**GACTACHVGGGTATCTAATCC |  |
| PCR for  *E. aedis* infection confirmation | FQEA187 | AGTGCGTACCGAGGCTATAAC | Duncan et al., 2012  (<doi.org/10.1002/ece3.199>) |
|  | RQEA310 | CTCAACGTTCATTGGGTAAGTTTC |  |
| qPCR to approximate bacterial load  (AAEL009496) | RpS7_F | TAGACACCCTGAAGTTGTTGCAAAT | MacLeod et al., 2021  (<https://doi.org/10.3389/fmicb.2021.645362>) |
|  | RpS7_R | TGTATATGCGCATTAGTCTCATCAA |  |
| qPCR to approximate bacterial load | 16S_F | TCCTACGGGAGGCAGCAGT |  |
|  | 16S_R | GGACTACCAGGGTATCTAATCCTGTT |  |
