## Supplementary figures and images for "The *Aedes aegypti* bacterial microbiota is robust to infection with the obligate microsporidian parasite *Edhazardia aedis*"

### Fig. S1

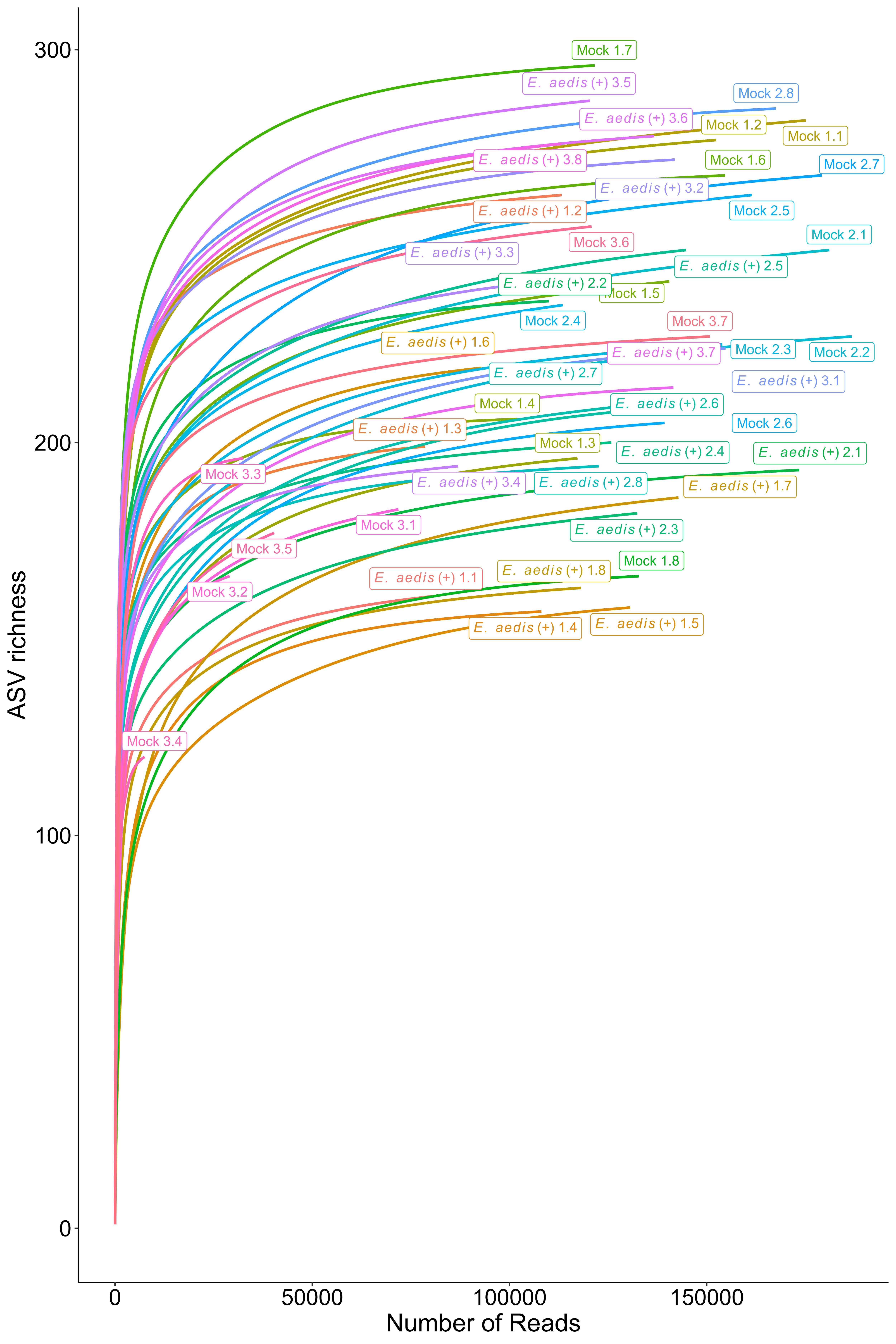

### Fig. S2

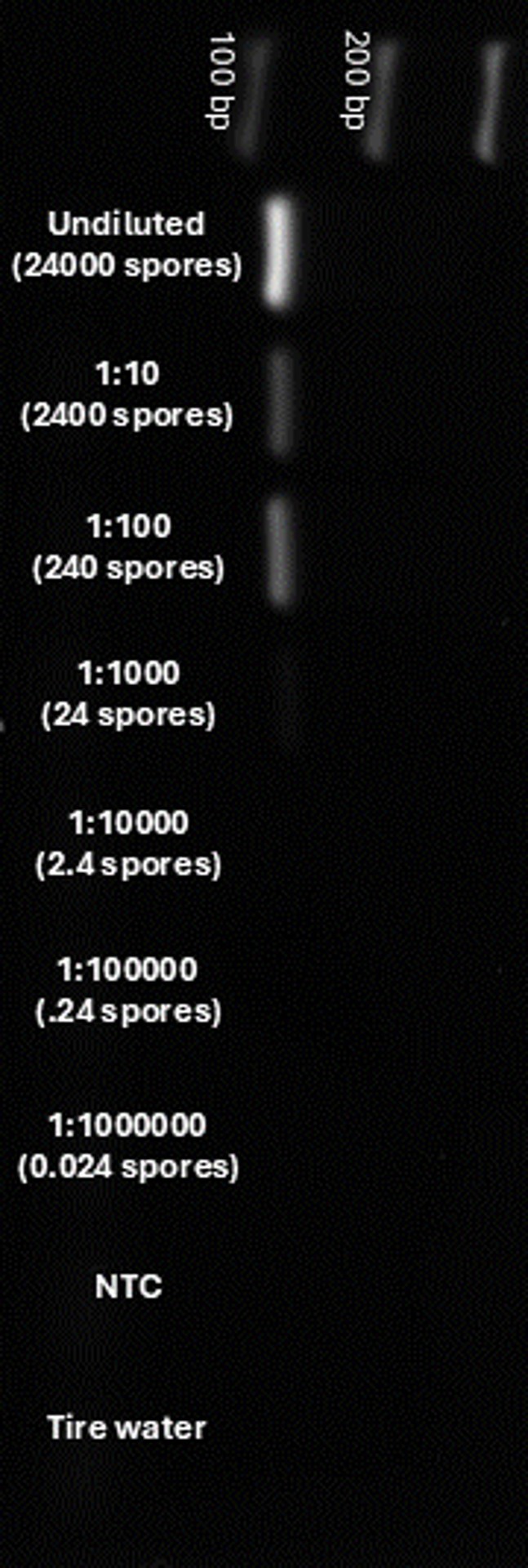

### Fig. S3

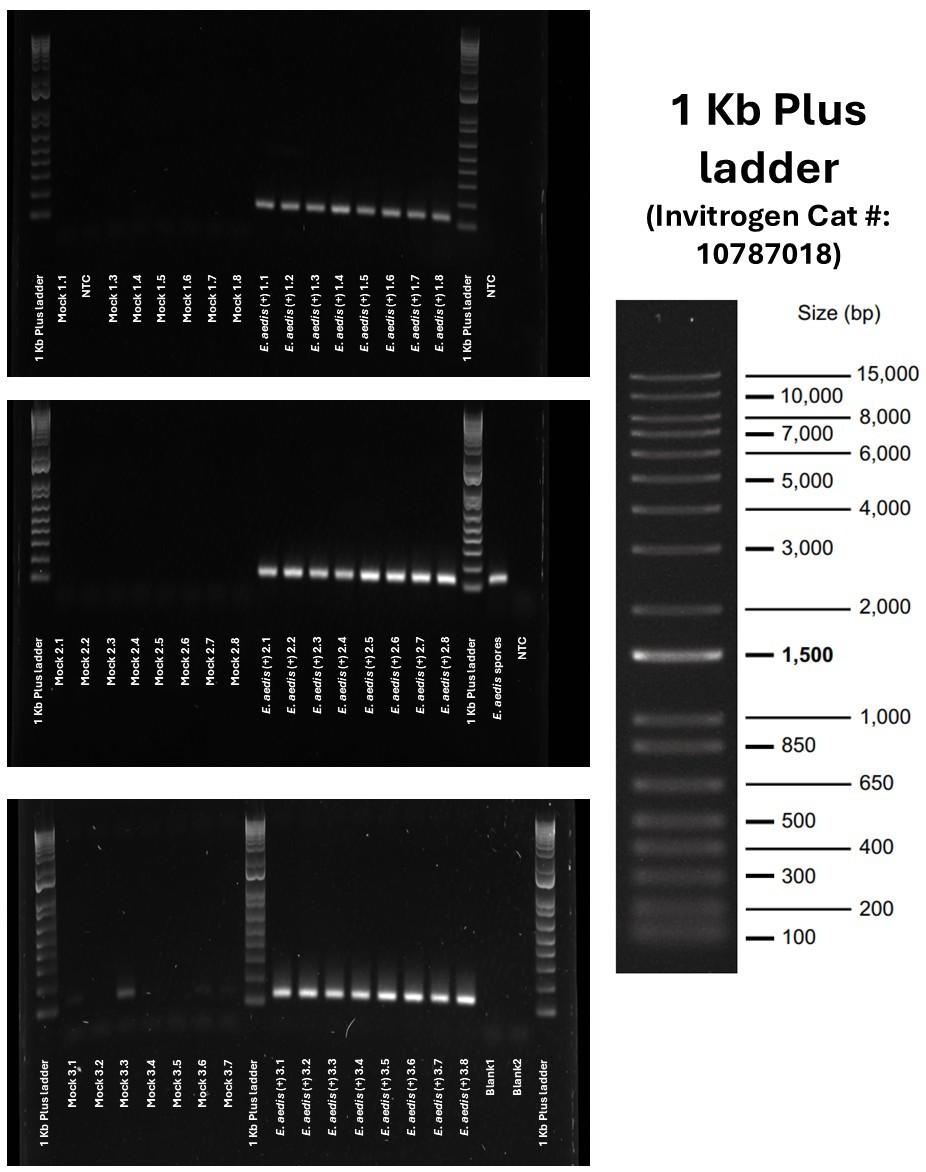

### Fig. S4

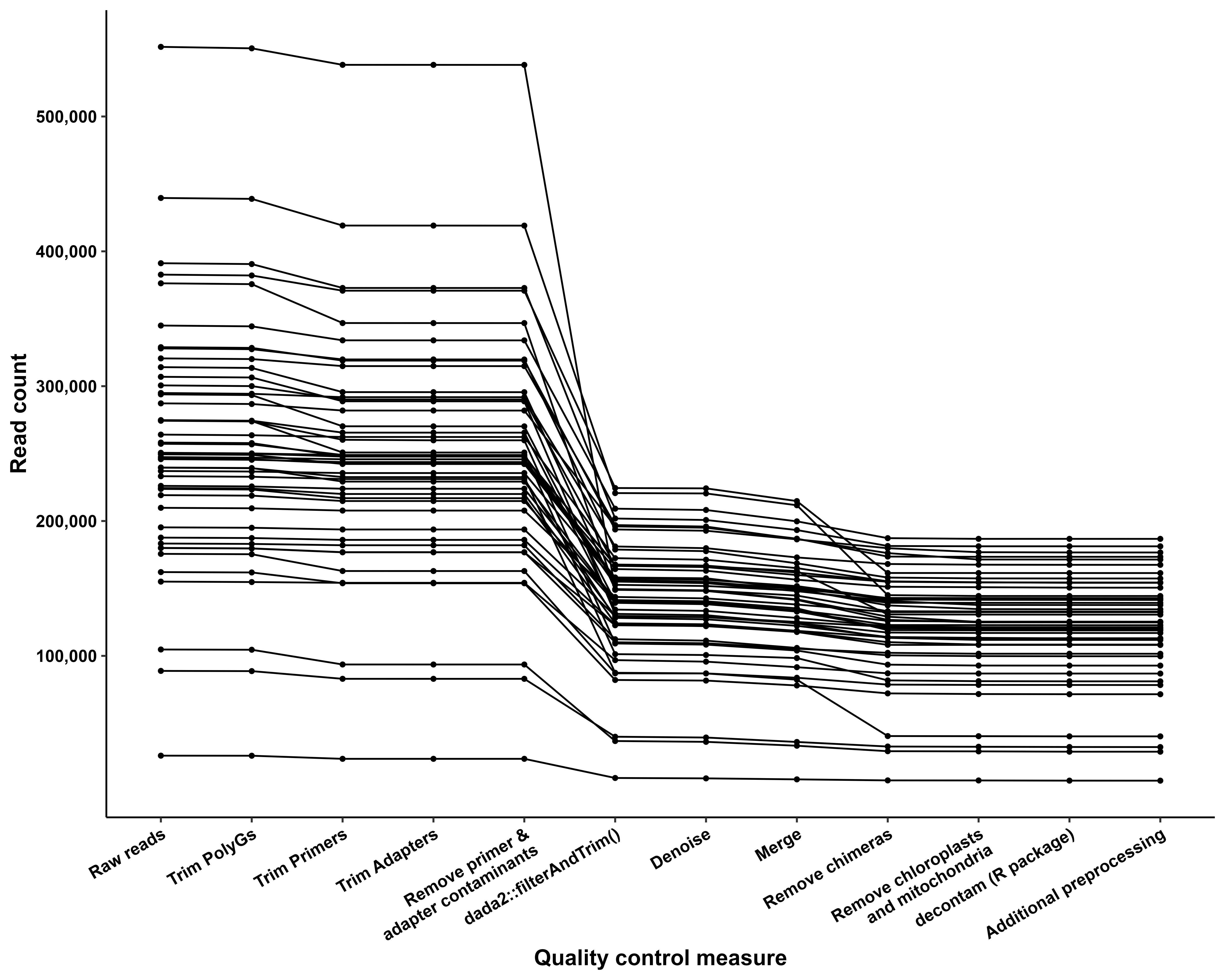
